# Genome-wide discovery of hidden genes mediating known drug-disease association using KDDANet

**DOI:** 10.1101/749762

**Authors:** Hua Yu, Lu Lu, Ming Chen, Chen Li, Jin Zhang

## Abstract

Many of genes mediating Known Drug-Disease Association (KDDA) are escaped from experimental detection. Identifying of these genes (hidden genes) is of great significance for understanding disease pathogenesis and guiding drug repurposing. Here, we presented a novel computational tool, called KDDANet, for systematic and accurate uncovering the hidden genes mediating KDDA from the perspective of genome-wide functional gene interaction network. KDDANet demonstrated the competitive performances in both sensitivity and specificity of identifying genes in mediating KDDA in comparison to the existing state-of-the-art methods. Case studies on Alzheimer’s disease (AD) and obesity uncovered the mechanistic relevance of KDDANet predictions. Furthermore, when applied with multiple types of cancer-omics datasets, KDDANet not only recapitulated known genes mediating KDDAs related to cancer, but also revealed novel candidates that offer new biological insights. Importantly, KDDANet can be used to discover the shared genes mediating multiple KDDAs. KDDANet can be accessed at http://www.kddanet.cn and the code can be freely downloaded at https://github.com/huayu1111/KDDANet.

## Introduction

The conventional development of novel promising drugs for treating specific diseases is a time-consuming and efforts-costing process, including discovery of new chemical entities, target detection and verification, preclinical and clinical trials and so on ^1^. Furthermore, the success rate for a new drug to be taken to market is very low, usually only 10% per year of drugs approved by FDA and thus prevents their use in actual therapy ^1^. This results in that pharmaceutical research faces a decreasing productivity in drug development and a sustaining gap between therapeutic needs and available treatments ^1^. Compared with traditional drug development, drug repositioning, i.e., finding the novel indications of existing drugs, offers the possibility for safer and faster drug development because of several steps of traditional drug development pipeline can be avoided during repurposing efforts ^2^. Many successful cases of repositioned drugs have been shown: from Minoxidil, designed for treatment of hypertension and now indicated for hair loss ^3^, to Sildenafil, developed for patients with heart problems and repurposed for erectile dysfunction ^4^. Yet, these examples are mainly based on clinical observations of secondary effects ^5^. Thanks to the advance in next generation omics sequencing and qualification technologies, a large volume of biomedical data, for example the pharmacogenomics datasets produced by Connective Map (CMap) project, The Cancer Genome Atlas (TCGA) project, Cancer Cell Line Encyclopedia (CCLE) project, Genomics of Drug Sensitivity in Cancer 1000 human cancer cell lines (GDSC1000) project and Library of Integrated Network-based Cellular Signatures (LINCS) project, has been rapidly accumulated for enabling drug repurposing ^6–14^. Based on these datasets, various computational methods have been designed for facilitating the process of drug repurposing (see Supplementary Note 1 for a mini review). To infer pharmacokinetic and pharmacodynamic drug-drug interactions and their associated recommendations, for example, Gottlieb et al. designed a similarity measure-based logistic regression classifier ^15^. For predicting drug side-effects, Tatonetti et al. presented an adaptive data-driven approach ^16^. To identify novel drug combinations, Zhao et al. integrated the molecular and pharmacological data and developed a novel computational approach ^17^. In addition, network-based method has also been employed to achieve the similar goal ^18^. Interestingly, Kuenzi et al. developed a deep-learning model of visible neural network, called DrugCell, to predict drug response and synergy in human cancer cells ^19^. For discovering novel drug indication, Gottlieb et al. developed PREDICT, which scored a possible drug-disease link by combining multiple drug-drug and disease-disease similarity measures ^20^. With the similar motivation and aim, an unsupervised and unbiased network-based proximity measure has also been designed ^21^. Moreover, Cheng et al. showed that the further integration of network proximity-based approach with large-scale patient-level longitudinal data can offer an effective platform for validating drug indications ^22^. For predicting drug-target interaction, Paolini et al. presented a global mapping of pharmacological space and probabilistic models by integrating multiple medicinal chemistry data ^23^. Different from the approach employed by Paolini et al., Campillos et al. used the phenotypic side-effect similarity to determine whether two drugs share a target ^24^. Besides, we and others also employed machine learning and network integration approaches for achieving the same goal ^25, 26^. To discover disease related genes, Gottlieb et al. designed “PRINCIPLE”, which employed classical network propagation algorithm ^27^. Wu et al. developed CIPHER that integrated human protein-protein interactions, disease phenotype similarities, and known gene-phenotype associations to capture the complex relationships between phenotypes and genotypes ^28^. Additionally, Ghiassian et al. identified disease gene modules based on systematic analysis of connectivity patterns of disease proteins in the human interactome ^29^. Furthermore, Zhou et al. and Menche et al. constructed disease-symptom and disease-disease relationship networks based on the biomedical literature databases and incomplete interactome, respectively ^30, 31^. Excitingly, Hofree et al. developed a new computational approach which employed network-based stratification (NBS) to uncover tumor subtypes by integrating somatic tumor genomes with gene networks^32^. Collectively, these methods have effectively exploited and integrated multi-level biomedical and omics data sources for understanding the pathology of diseases and mechanisms of drug actions and thus accelerated drug repurposing.

Theoretically, drug repurposing has been proposed based on two molecular aspects. I) On one hand, complex diseases often involve multiple genetic and environmental determinants, including multi-factor driven alterations and dysregulation of a series of genes ^33, 34^, which will propagate and perturb certain biological processes by the interactions among molecules, leading to the onset of diseases. II) On the other hand, one drug can exert impacts on many targets and perturb multiple biological processes ^34, 35^. As a result, the genes of shared biological pathways in the cellular network manifested in certain disease state and induced by a known drug administration suggest potential drug repurposing ^33–38^. However, many of these genes mediating Known Drug-Disease Associations (KDDAs) across various types of diseases have not yet been identified (see the Results section and Supplementary Figure S1a-S1b for supporting data of this claim). Therefore, developing the appropriate theoretical computational tools from the perspective of molecular interaction network to unveil the KDDA genes missed from experiments (hidden genes) is of great significance for understanding disease pathogenesis and guiding drug repurposing. The network-based computational methods have been designed for facilitating drug repurposing which linked drugs to targets or connected diseases to genes or associated drugs with diseases ^18, 21, 22, 26–29, 39^. Nevertheless, the publicly available computational tools specially tailored for simultaneously bridging drugs, genes, and diseases have not been fully developed. To our knowledge, some computational tools have been designed to identify drug-gene-disease co-module. Kutalik et al. and Chen et al. developed Ping-Pong Algorithm (PPA) and Sparse Network-regularized Partial Least Square (SNPLS) to identify co-modules related to specific cancer cell lines of NCI-60 and Cancer Genome Projects, respectively ^40, 41^. However, these two methods need to integrate gene-expression and drug-response data of cancer cell lines for constructing models and do not carry out prediction for other types of diseases. The other two methods of comCIPHER and DGPsubNet have been designed to identify the coherent subnetworks linking drugs and diseases (not limited in cancer) with the related genes ^42, 43^. The common shortcomings of these methods are the identified co-modules including multiple drugs and diseases and thus are unable to uncover genes specifically mediating individual KDDA. To this end, we designed a novel computational pipeline, called KDDANet, which uses known functional gene interaction network to identify hidden genes of cellular pathways mediating KDDA in a genome-wide scale. Our KDDANet pipeline depends on three existing network algorithms: minimum cost network flow optimization, depth first searching and graph clustering algorithm. The minimum cost network flow optimization has been effectively employed to identify cellular response subnetwork connecting genetic hits and differentially expressed genes, including components of the response that are otherwise hidden or missed from experiments ^44^. KDDANet can be applied to two contexts: 1) uncovering hidden genes mediating the association between a query drug and its related disease (SDrTDi); 2) unveiling hidden genes mediating the association between a query disease and its related drug (SDiTDr). The computational procedure of KDDANet in SDrTDi context was showed in Fig. 1 (see Method for details). KDDANet first built a unified flow model by integrating query drug, genes, and all related diseases into a heterogeneous network (Fig. 1a). Then, the minimum cost flow optimization was designed and implemented to identify gene subnetwork mediating the association between the query drug to all its related diseases (Fig. 1b). Finally, depth first searching, and Markov clustering algorithm were adopted to further uncover gene modules mediating the association between the query drug and each its related disease (Fig. 1c-1e). The outputs of KDDANet were validated against existing literature and multi-omics’ datasets (Fig. 1f). To apply KDDANet in SDiTDr context, what the user need was just to simply rebuilt the unified flow network model (see Methods for details).

**Fig. 1:**
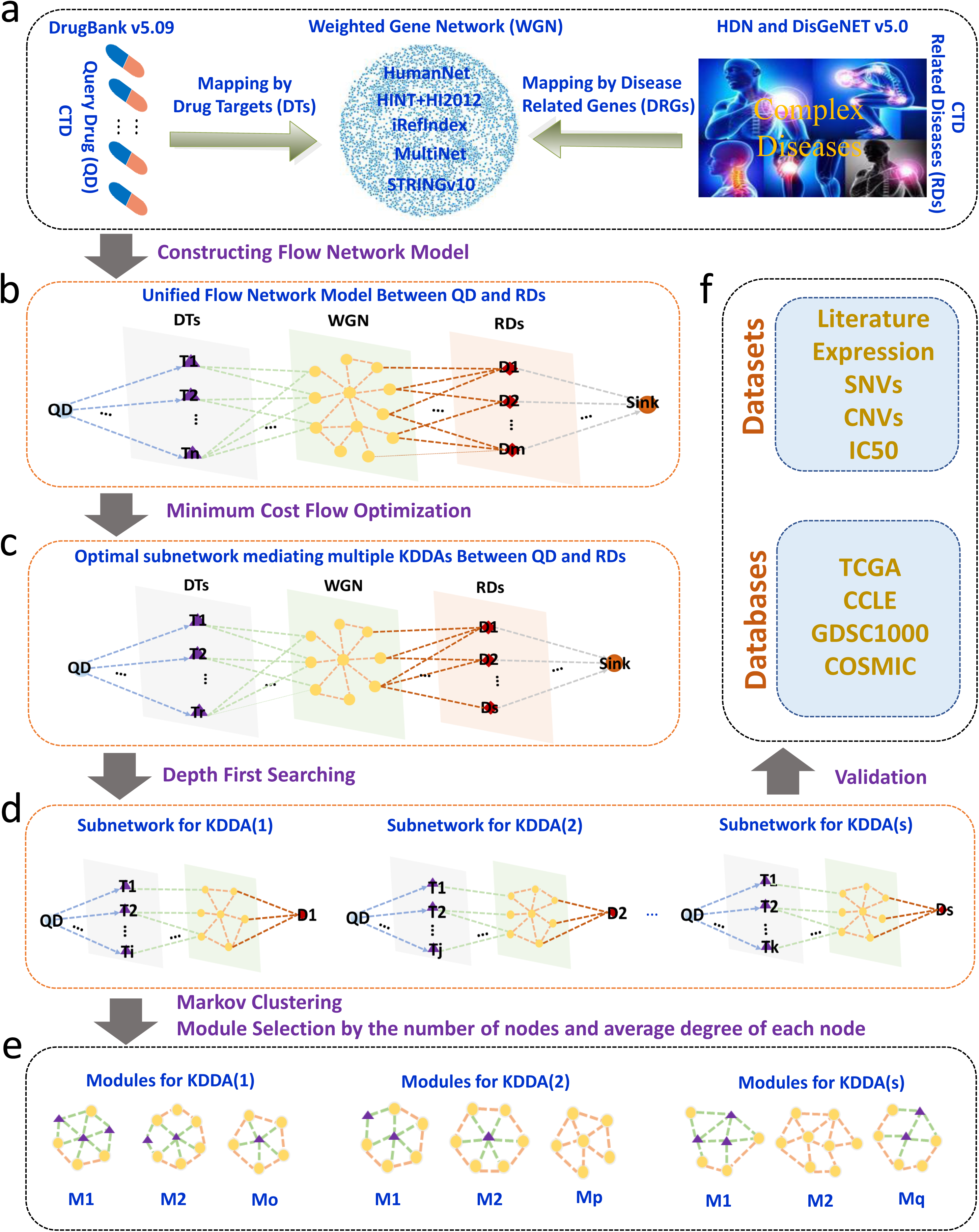
Schematic illustration of KDDANet computational pipeline in SDrTDi context. **a)** Mapping query drug (QD) and all its related diseases (RDs) into the weighted gene network (WGN) through known drug-target relationships and gene-disease associations. **b)** Constructing a unified flow network model for each QD and RDs (multiple KDDAs). **c)** Identifying the highest probability gene subnetwork mediating multiple KDDAs by minimum cost flow optimization. **d)** Identifying gene subnetwork mediating individual KDDA by depth first searching. **e)** Identifying gene interaction modules mediating individual KDDA by Markov clustering (MCL). **f)** Validating the prediction results of KDDANet by existing knowledgebases.

The key novelty of the KDDANet method lay in that the original designation of a unified flow network model and effective implementing multiple network-based algorithms on this unified flow network model. We demonstrated that KDDANet showed competitive performance in discovering known and novel genes mediating KDDA through a comparison with existing methods. KEGG pathway enrichment analysis on KDDANet resulting subnetworks and case studies on Alzheimer’s disease (AD) and obesity further showed the mechanistic relevance of KDDANet predictions. Validated with multiple types of cancer-omics’ datasets, KDDANet did not only revealed known genes mediating KDDAs associating drug with cancer, but also uncovered new candidates that offer novel biological insights. Particularly, our results demonstrated that KDDANet can reveal the shared genes mediating multiple KDDAs. These outcomes showed that importance of incorporating hidden genes in drug discovery pipelines. For facilitating biomedical researchers to explore the molecular mechanism of KDDA and guiding drug repurposing, an online web server, http://www.kddanet.cn, was provided for user to access the subnetwork of genes mediating KDDA, and the source codes of KDDANet were freely available at https://github.com/huayu1111/KDDANet. In summary, we developed an effective and universal computational tool and an online web source for accurate and systematic discovering hidden genes mediating KDDA and thus providing novel insights into mechanism basis of drug repurposing and disease treatments. We believed that KDDANet can provide additional contributions to the development of new therapies.

## Results

### Evaluation of the performance and general applicability of KDDANet method

We first checked whether the potential genes mediating KDDA across various types of diseases have been experimentally identified. For a given KDDA, we proposed a hypothesis that the Known Drug Target Genes (KDTGs) and Known Disease Related Genes (KDRGs) should be highly overlapped if the genes mediating this KDDA have been fully identified. We analyzed the overlap between Known Drug Target Genes (KDTGs) and Known Disease Related Genes (KDRGs) of 53124 KDDAs obtained from Comparative Toxicogenomics Database (CTD) ^12^. For a KDDA, we defined the overlap ratio as the number of shared genes between KDTGs and KDRGs divided by the number of total KDTG and KDRG genes. We observed that the most gene sets demonstrated extremely low overlap ratio, with the increasing of gene number, the overlap ratio was sharply decreased (Supplementary Figure 1a). We then checked the overlap ratio in different types of diseases and found the overlaps between KDTGs and KDRGs were small for all 19 disease types in our dataset (Supplementary Figure 1b). The discrepancy between KDTGs and KDRGs indicated that each gene set alone provided only a limited and biased view of KDDA, many of true genes in the cellular pathways mediating KDDA were not identified from the experiments but otherwise hidden. To address this, we designed a novel computational tool, KDDANet, which effectively integrated minimum cost flow optimization, combined with depth first searching and graph clustering to systematically discover the hidden genes of cellular pathways mediating KDDA (see Methods for details).

To examine whether KDDANet can capture true genes mediating KDDA, we introduced two new concepts: “known true KDDA genes” (KTKGs) and “novel true KDDA genes” (NTKGs). For a given KDDA, KTKGs was defined as the shared genes between KDTGs and KDRGs inputted for constructing KDDANet flow network model (see Dataset for details). To obtain NTKGs, we collected a set of drug’s non-target genes (A) from SMPDB 2.0 database ^45^ that were included in the drug’s ADME pathways and were responsible for mediating KDDA. Meanwhile, we collected a recently updated set of disease-related genes (B) from DisGeNet v6.0 database ^46^, which were not included in KDDANet flow network model. Based on these, we defined, for each KDDA, the NTKGs as the shared genes between A and B. A parameter γ effected the size and quality of KDDANet output subnetwork, higher γ values will identify more gene links mediating KDDA but with lower confidence. Using gene set enrichment analysis, we observed that KDDANet can consistently and effectively capture the KTKGs and NTKGs mediating KDDA under different γ settings (see Supplementary Note 2, Supplementary Note 3, and Supplementary Figure 1c-1l for details).

Based on this, we compiled a standard set of positive and negative KDDA genes for each KDDA to unbiasedly evaluate the capability of KDDANet method on uncovering the true genes mediating KDDA using Receiver Operating Characteristic (ROC) and Precision-Recall (PR) curves (see Methods for details). We observed that the performance of KDDANet was obviously better than random permutation across a widely settings of *y* (Fig. 2a and Fig. 2b). We next selected γ = 6 for SDrTDi and γ = 8 for SDiTDr for subsequent evaluation (see Supplementary Note 4 for reasons). With this setting, we further compared KDDANet with other existing methods for drug-gene-disease co-module discovery, including SNPLS ^40^, PPA ^41^, comCHIPER ^42^ and DGPsubNet ^43^ (see Methods for the details of performance comparison). Among all the methods tested, KDDANet demonstrated the competitive performance with the averages of AUROC 0.733 and 0.715 and AUPRC 0.793 and 0.825 in SDrTDi and SDiTDr contexts, respectively (Fig. 2c and Table 1). These results suggested that KDDANet was an effective tool for achieving the goal of uncovering true genes mediating KDDA genome wide. We next tested whether KDDANet had the general application value across a variety of disease types and found that KDDANet can make effective capturing of true genes mediating KDDA for all 19 types of diseases (Supplementary Figure 2a-2d). We further tested KDDANet using different types of networks, including HINT+HI2012, iRefIndex, MultiNet and STRINGv10 ^94^. We found that the performances of KDDANet were consistent well across all these networks (Fig. 2d, Supplementary Figure 2e and Supplementary Figure 2f). Collectively, these results fully demonstrated that KDDANet was an effective and general computational tool for uncovering hidden genes mediating KDDA across broad types of diseases.

**Fig. 2:**
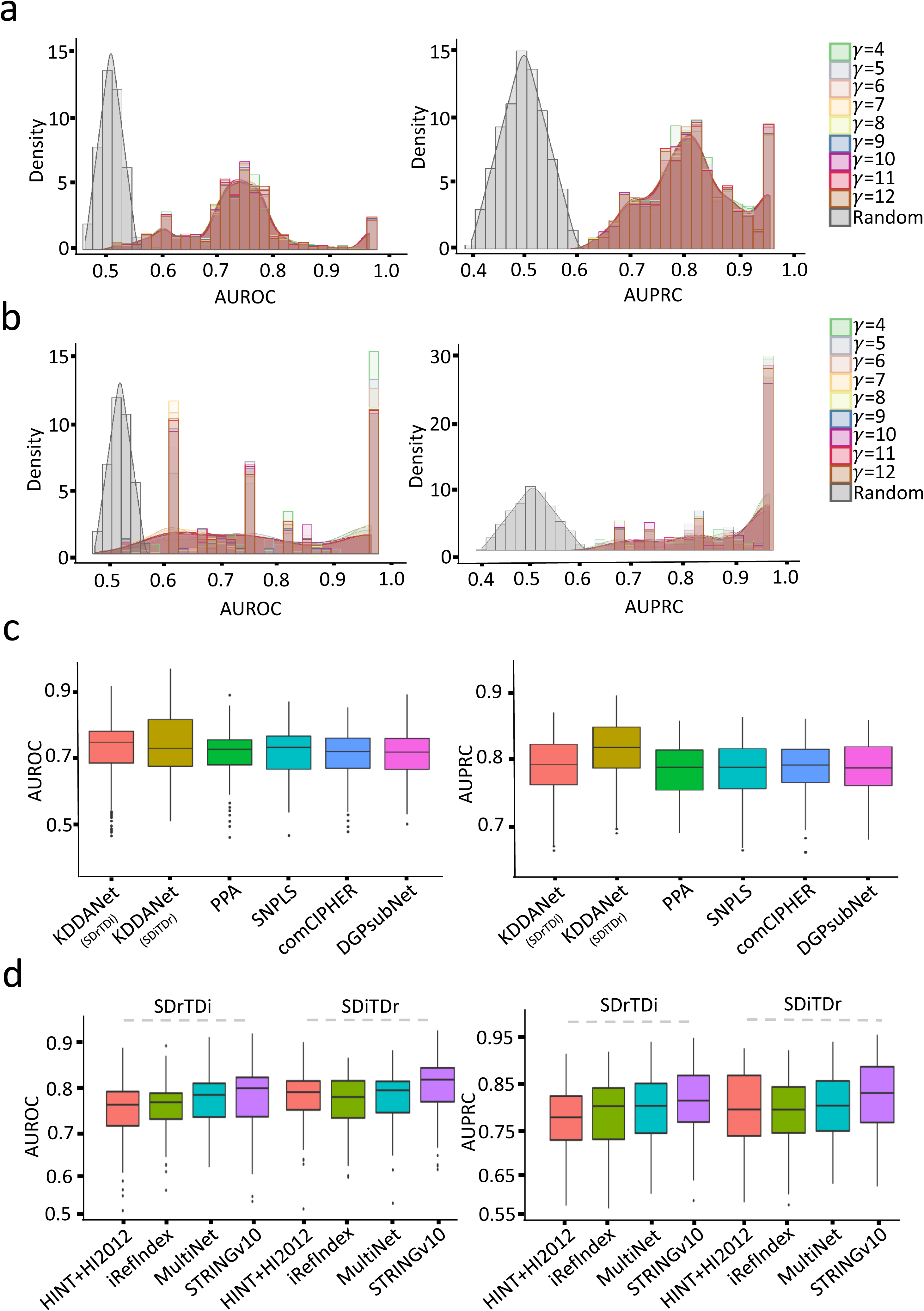
Performance evaluation of KDDANet method. **a)** The density curve of AUROC and AUPRC of KDDANet with different γ setting and permutation test in SDrTDi context. **b)** The density curve of AUROC and AUPRC of KDDANet with different γ setting and permutation test in SDiTDr context. **c)** Comparison of AUROC and AUPRC of KDDANet with PPA, SNPLS, comCHIPER and DGPsubNet. **d)** AUROC and AUPRC of KDDANet with different types of gene interaction networks.

**Table 1.** Average AUC values of different computational tools for identifying hidden genes mediating KDDA.

| Method | KDDANet(SDrTDi) | KDDANet(SDiTDr) | SNPLS | PPA | comCHIPER | DGPsubNet |
| --- | --- | --- | --- | --- | --- | --- |
| Average AUROC | 0.733 | 0.715 | 0.712 | 0.713 | 0.703 | 0.706 |
| Average AUPRC | 0.793 | 0.825 | 0.775 | 0.776 | 0.778 | 0.777 |

### Mechanistic relevance of KDDANet predictions

We examined whether the enriched pathways of KDDANet resulting subnetworks have the mechanistic relevance with KDDA by carrying out a global enrichment of all predicted KDDA subnetwork genes against 53 classical KEGG pathways. The obtained enrichment results can be validated by existing knowledge (see Supplementary Note 5 and Supplementary Figure 3a-3b for details). We aimed to provide two cases to intuitively describe the mechanistic relevance of KDDANet resulting subnetwork. Phylloquinone (DB01022)-Alzheimer’s disease (AD) (104300) association has been reported in previous study ^47^. A subnetwork including 46 genes and 44 links were uncovered mediating this association (Fig. 3a). Interestingly, for this subnetwork, two separated gene modules (M1 and M2) were detected (Fig. 3a). The AUROC and AUPRC values of for this KDDANet resulting subnetwork were 0.864 and 0.795, respectively (Supplementary Figure 3c). Two known targets of phylloquinone and 18 AD-related genes were identified in this subnetwork. Against with genome background, this subnetwork captured 4 NTKGs with ∼86-fold enrichment and adjusted *p*-value of 9.716e-09 (Hypergeometric test and Bonferroni correction). The top 10 enriched KEGG terms of this subnetwork, such as Phospholipase D signaling pathway and Neurotrophin signaling pathway ^48, 49^, were closely related with AD (Fig. 3b). The module M1 mainly functioned in Insulin signaling pathway, ErbB signaling pathway, FoxO signaling pathway and growth hormone synthesis, secretion, and action, which played important roles in neural system development and the onset and development of AD ^50–52^ (Supplementary Figure 3d). The enriched KEGG pathways of M2 genes, including Complement and coagulation cascades, Glycolysis and AGE-RAGE signaling pathway in diabetic complications, were dysfunction in AD ^53–55^ (Supplementary Figure 3d). We further analyzed the published RNA-seq data to detect the expressional change of these two modules in normal individuals and AD patients ^56^. We observed that the averaged expression level of M1 genes was significantly upregulated in AD (Supplementary Figure 3e). Interestingly, a predicted novel gene, STAT3, was obviously activated in AD patients (Fig. 3c). Two newest studies published in years 2019 and 2020 reported that STAT3 was a potential therapeutic target for cognitive impairment in AD ^57, 58^. Another predicted novel gene, GNAI1, was also obviously activated in AD patients. This gene was contained in the causal pathways associated with an imaging endophenotype characteristic of longitudinal structural change in the brains of patients with AD ^59^. For module M2 genes, the obviously expressional changes between normal individuals and AD patients were not observed (Supplementary Figure 3e). However, we found that a predicted novel KDDA gene, GC, was activated in AD patients (Fig. 3c). This gene encoded Vitamin D Binding Protein, which was recently evidenced as a potential therapeutic agent for the treatment of AD ^60^.

**Fig. 3:**
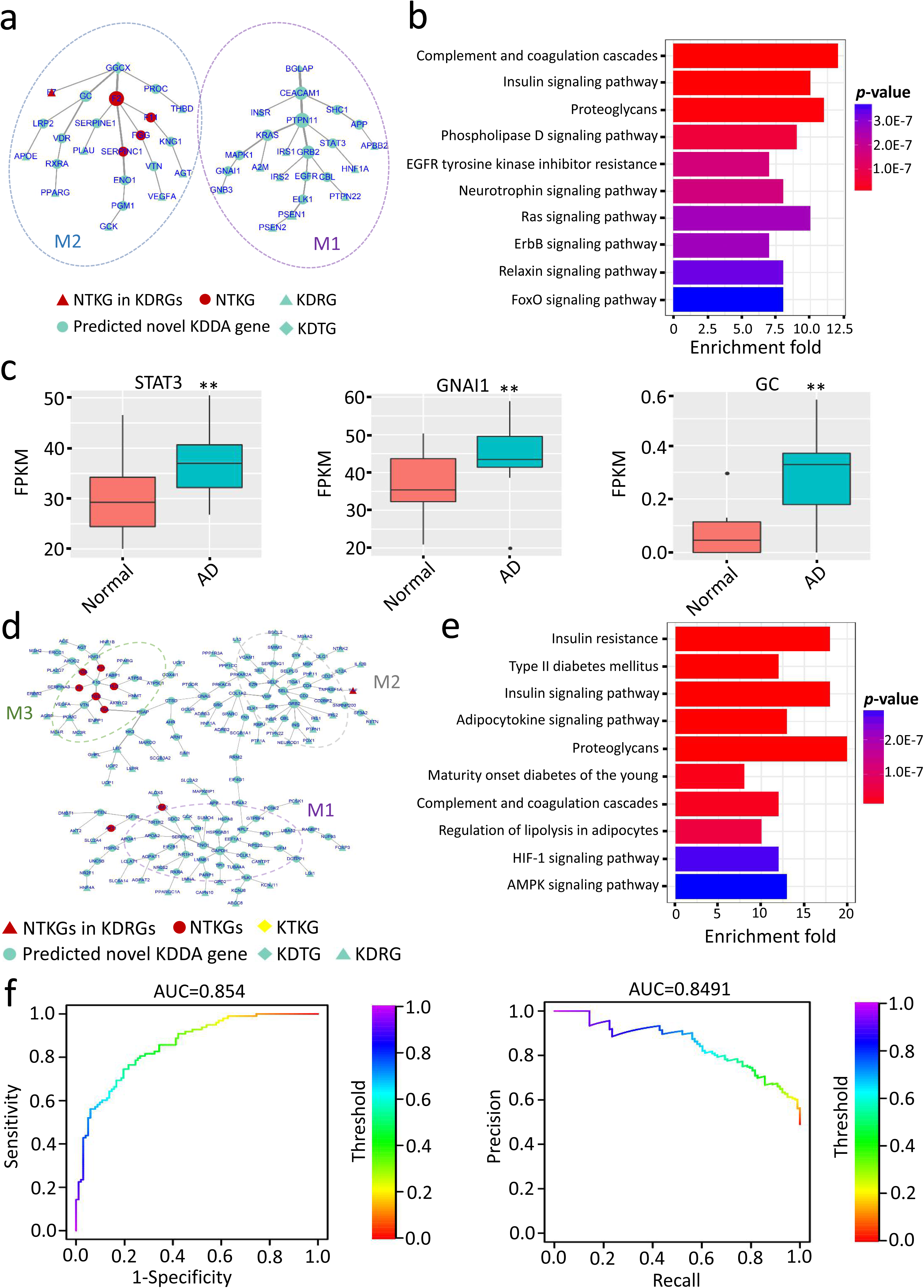
Mechanistic relevance of KDDANet prediction results. **a)** KDDANet resulting subnetwork mediating phylloquinone (DB01022)-Alzheimer’s disease (AD, 104300) association. **b)** Top 10 enriched KEGG terms of subnetwork genes mediating phylloquinone-AD association. **c)** Expression level of STAT3, GNAI1 and GC in normal individuals and AD patients, ** *p*-value < 0.01, calculated by Mann-Whitney U test. **d)** KDDANet resulting subnetwork for heparin (DB01109)-obesity (601665) association. **e)** Top 10 enriched KEGG terms of subnetwork genes mediating heparin-obesity association. **f)** ROC and PR curves of KDDANet gene subnetwork mediating heparin-obesity association. In the subnetworks, the size of a gene node was proportional to its network degree; The thickness of a network edge was proportional to its flow amount. Fragments Per Kilobase Of Exon Per Million Fragments Mapped, FPKM.

The association between heparin (DB01109) and obesity (601665) was inferred by multiple genes as described in CTD database. For this association, KDDANet predicted a subnetwork containing 168 edges connecting 169 genes (Fig. 3d). We found that two known drug target genes and 67 disease-related genes were captured in this subnetwork. Particularly, 8 genes were the NTKGs with ∼52-fold enrichment and adjusted *p*-value of 1.485e-10 (Hypergeometric test and Bonferroni correction). Three genes used to infer this KDDA, including ARK1, PARP1 and TNF, were also effectively captured in the resulting subnetwork. The top 10 enriched functions of this subnetwork were showed in Fig. 3e. As expected, Insulin resistance, Type II diabetes mellitus, Insulin signaling pathway and Adipocytokine signaling pathway were the frequently reported events and molecular processes associated with obesity ^61, 62^. In addition, Proteoglycans, Lipolysis and AMPK signaling pathway were also highly related with Insulin resistance ^63–65^. In consistent with this, the AUROC and AUPRC values of KDDANet for this subnetwork were 0.854 and 0.849, respectively (Supplementary Figure 3e). We next delineated the subnetwork into gene modules. The enriched functions of top 3 gene modules were demonstrated in Supplementary Figure 3f. The genes of module M1 mainly participated in Glycolysis and Carbon metabolism. Further analysis of public RNA-seq data of normal individuals and obesity patients ^66, 67^ demonstrated the significantly repressed expression of M1 genes in patients with obesity, such as GADPH (Supplementary Figure 3g and 3h). This was consistent with the fact that enhancing the level of glycolysis reduced obesity ^68^. The mainly enriched pathways of M2 genes were related to Insulin resistance, a frequently happened event in obesity patients. Interestingly, Cell adhesion molecules was a significantly enriched KEGG term of M2 genes (Supplementary Figure 3f), which was elevated in patients with obesity ^69^. The M3 genes functioned in Complement and coagulation cascades and Platelet activation (Supplementary Figure 3f). As reported, these two terms were closely associated with obesity ^70, 71^. In support with these, the expression of genes in M2 and M3 were activated in patients with obesity (Supplementary Figure 3g and Supplementary Figure 3h). Together, these results indicated that KDDANet can serve as a useful tool to unveil the molecular basis of KDDA.

### KDDANet provided novel molecular insights on KDDAs related to cancer

Cancer was a frequently happened complex genetic disease caused by DNA abnormalities ^72^. For this reason, substantial genetic, genomic and pharmacogenomics efforts, including TCGA, CCLE and GDSC1000 ^7–9^, have been undertaken to improve existing therapies or to guide early-phase clinical trials of compounds under development. With these efforts, an increasing amount of available high-throughput data sets at both levels of genomic data and pharmacogenomics data were produced at recent years. In addition, COSMIC, the world’s largest and most comprehensive resource for exploring the impact of somatic mutations in human cancer, collected a catalogue of genes with mutations that were causally implicated in cancer (https://cancer.sanger.ac.uk/cosmic). With these datasets, we observed that the genes of KDDANet resulting subnetworks mediating the associations between drugs and cancer were significantly enriched in COSMIC Cancer Gene Census, and these genes harbored more oncogenic alterations in tumor samples than randomly selected genes (Supplementary Note 6, Supplementary Figure 4a and Supplementary Figure 4b). Moreover, we found that oncogenic alterations of genes in KDDANet resulting subnetworks mediating the associations between drugs and cancer were more correlated with the responses of cancer cell lines under anti-cancer drug treatment than randomly selected genes (see Supplementary Note 5, Supplementary Figure 4c-4f). We provided two detailed examples to describe the potential values of KDDANet in revealing novel genes mediating the associations between drugs and cancer.

Sotalol (DB00489) was normally used to treat life threatening ventricular arrhytmias. It has been reported that sotalol was associated with decreased prostate cancer (176807) risk ^73^. For sotalol-prostate cancer association, KDDANet predicted a subnetwork consisting of 31 genes and 28 links (Fig. 4a). By applying MCL with default parameters, this subnetwork was further decomposed into three gene modules, M1, M2 and M3. All three known target genes of sotalol were included in this subnetwork. Meanwhile, this subnetwork also captured 12 prostate cancer-related genes with a novel NTKG of PRKACA. The top 10 enriched KEGG terms of this subnetwork were showed in Fig. 4b. Among these, PI3K-Akt signaling pathway, MAPK signaling pathway and FoxO signaling pathway were related to cancer formation and development. The relationships between EGFR tyrosine kinase inhibitor resistance, Relaxin signaling pathway, AGE-RAGE signaling pathway and prostate cancer have been widely investigated and reported in the previous works ^74–76^. In addition, Autophagy and Focal adhesion were two widely observed processes in cancer ^77, 78^. As expected, the enriched KEGG signaling pathways of M1 were closely related to tumorigenesis (Supplementary Figure 4g). The expression of M1 genes were activated in tumor samples, such as an oncogene YWHAE (Supplementary Figure 4h and Supplementary Figure 4i). Interestingly, M1 captured that IGF1R, a gene encoding insulin-like growth factor receptor, harbored SNVs and CNVs in TCGA prostate cancer samples, and has been reported to be oncogenic genes of prostate cancer ^79^. The genes of M2 were not enrich to any KEGG term and but have lower expression in TCGA tumor samples (Supplementary Figure 4h). Surprisingly, SPARC, a reported prostate cancer-related gene ^80^ with significantly lower expression in TCGA tumor samples was captured in this module (Supplementary Figure 4i). Moreover, we found that both SPARC and COL1A1 harbor SNVs and CNVs in TCGA prostate cancer samples and their expressions were obviously correlated to the survival of patients (Fig. 4c). The M3 genes mainly participated in ErbB signaling pathways, a biological process involved in prostate cancer progression ^81^. The expression of M3 genes were not obviously changed in TCGA prostate cancer samples (Supplementary Figure 4h). However, we found that GRB2 was over-expressed in TCGA prostate cancer samples and obviously correlated with the survival of patients at the later stage of disease (Supplementary Figure 4i and Fig. 4c).

**Fig. 4:**
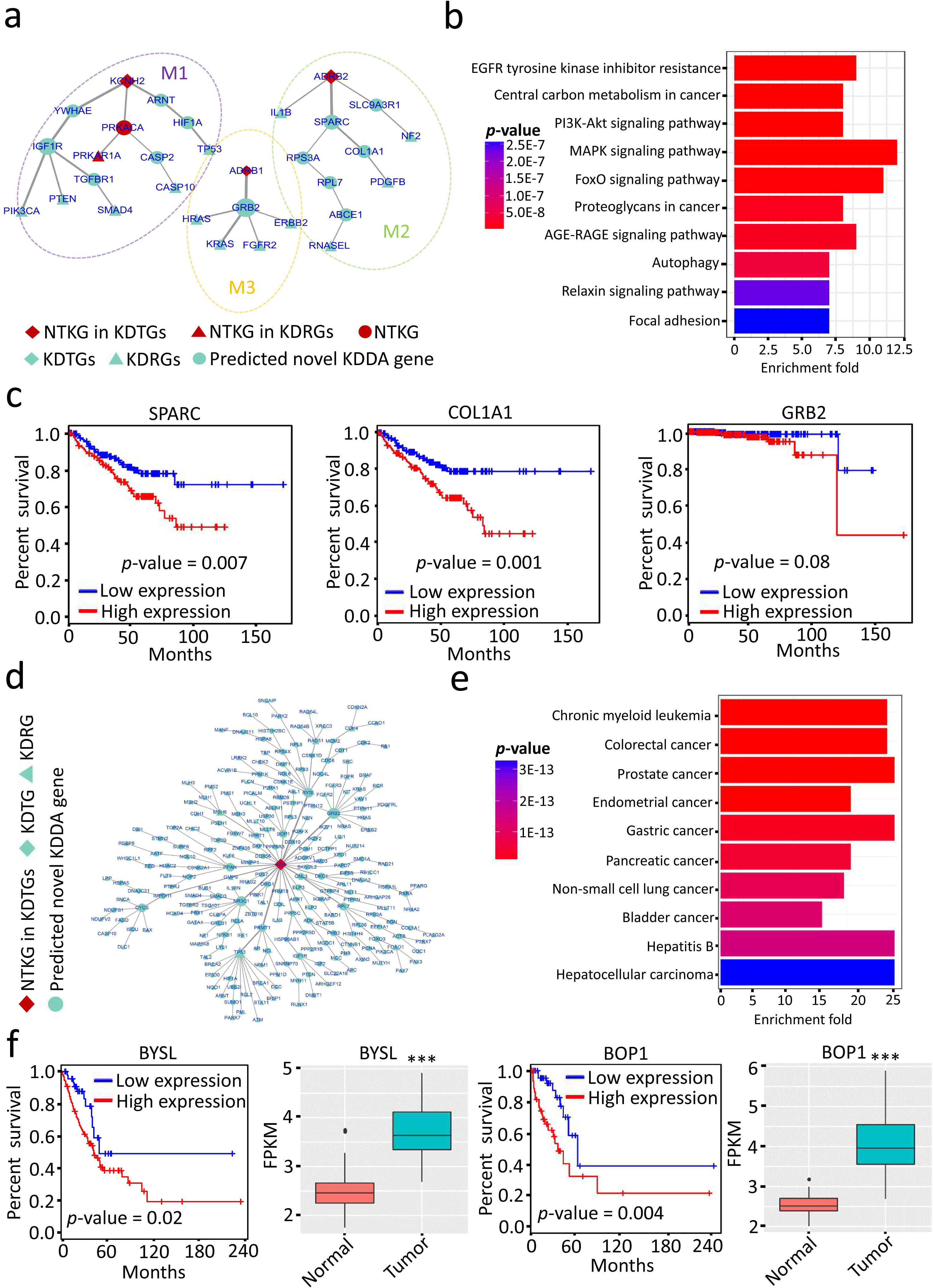
KDDANet provided novel molecular insights on KDDAs related to cancer. **a)** KDDANet resulting subnetwork mediating sotalol (DB00489)-prostate cancer (176807) association. **b)** Top 10 enriched KEGG terms of subnetwork genes mediating sotalol-prostate cancer association. **c)** Gene expression-based survival analysis for CASP2, SPARC and GRB2 in patients with prostate cancer, obtained by GEPIA online analysis (http://gepia.cancer-pku.cn/about.html). **d)** KDDANet gene subnetwork mediating nebularine (DB04440)-lung cancer (211980) association. **e)** Top 10 enriched KEGG terms of subnetwork genes mediating nebularine-lung cancer association. **f)** Gene expression-based survival analysis for BYSL and BOP1 in patients with TCGA LUAD lung cancer (obtained by GEPIA online analysis (http://gepia.cancer-pku.cn/about.html) and their expression levels in lung cancer tumor samples and adjacent normal tissue samples, *** *p*-value < 0.001, calculated by Wilcox signed rank test. In the subnetworks, the size of a gene node was proportional to its network degree; The thickness of a network edge was proportional to its flow amount.

Another example was the association between nebularine (DB04440)-lung cancer (211980) that was inferred by ADA targeted by nebularine ^82^. For this association, KDDANet predicted 237 genes connected by 238 links that constituted a subnetwork without apparent modular structure (Fig. 4d). The only known target gene ADA of nebularine was connected to 119 lung cancer-related genes and 117 predicted novel KDDA genes in this subnetwork. As expected, KEGG enrichment demonstrated that the genes in this subnetwork were involved in various cancers (Fig. 4e). Interestingly, we found that two highly connected novel genes BYSL and BOP1 were significantly over-expressed in TCGA lung cancer tumor samples and their expression levels were obviously correlated with the survival of lung cancer patients (Fig. 4f). These results indicated that KDDANet not only captured known genes mediating KDDAs linking drug with cancer, but also uncovered novel candidates that offered novel biological insights.

### KDDANet uncovered the shared genes mediating multiple KDDAs

Comprehensive analysis above fully demonstrated that KDDANet can uncover true genes mediating individual KDDA. We further asked whether KDDANet can reveal the shared genes mediating multiple KDDAs. We answered this from two aspects as follow: I) Multiple Diseases associating with One Drug (MDiODr); II) Multiple Drugs associating with One Disease (MDrODi). Considering the practical merits for the first one analysis, we required multiple diseases belongs to the same type of diseases. To evaluate the capability of KDDANet for revealing the shared genes mediating multiple KDDAs, we produced meta-subnetworks by integrating multiple KDDANet resulting subnetworks for 12386 MDiODr combinations and 773 MDrODi combinations produced in SDrTDi context, and 12189 MDiODr combinations and 773 MDrODi combinations in SDiTDr context. The weight of an edge in the meta-subnetwork was defined as the number of KDDA resulting subnetworks containing this edge divided by the total number of KDDA resulting subnetworks. The higher weight value of a link in the meta-subnetwork indicated more conservation and commonality. Thus, we used the weight value to evaluate the capacity of KDDANet in unveiling the shared gene interactions mediating multiple KDDAs. We carried out a permutation test by producing random meta-subnetworks with the same number of edges for comparing with random background. As shown in Fig. 5a, Supplementary Figure 5a and Supplementary Figure 5b, the weights of KDDANet meta-subnetworks were significantly higher than random meta-subnetworks across different types of diseases in SDrTDi context. This indicate that the genes tend to be shared in KDDANet meta-subnetworks than random one. We also conducted the same analysis in SDiTDr context and obtained the similar results (Fig. 5b, Supplementary Figure 5c and Supplementary Figure 5d). Collectively, these results indicated that KDDANet can effectively uncover the shared genes mediating multiple KDDAs.

**Fig. 5:**
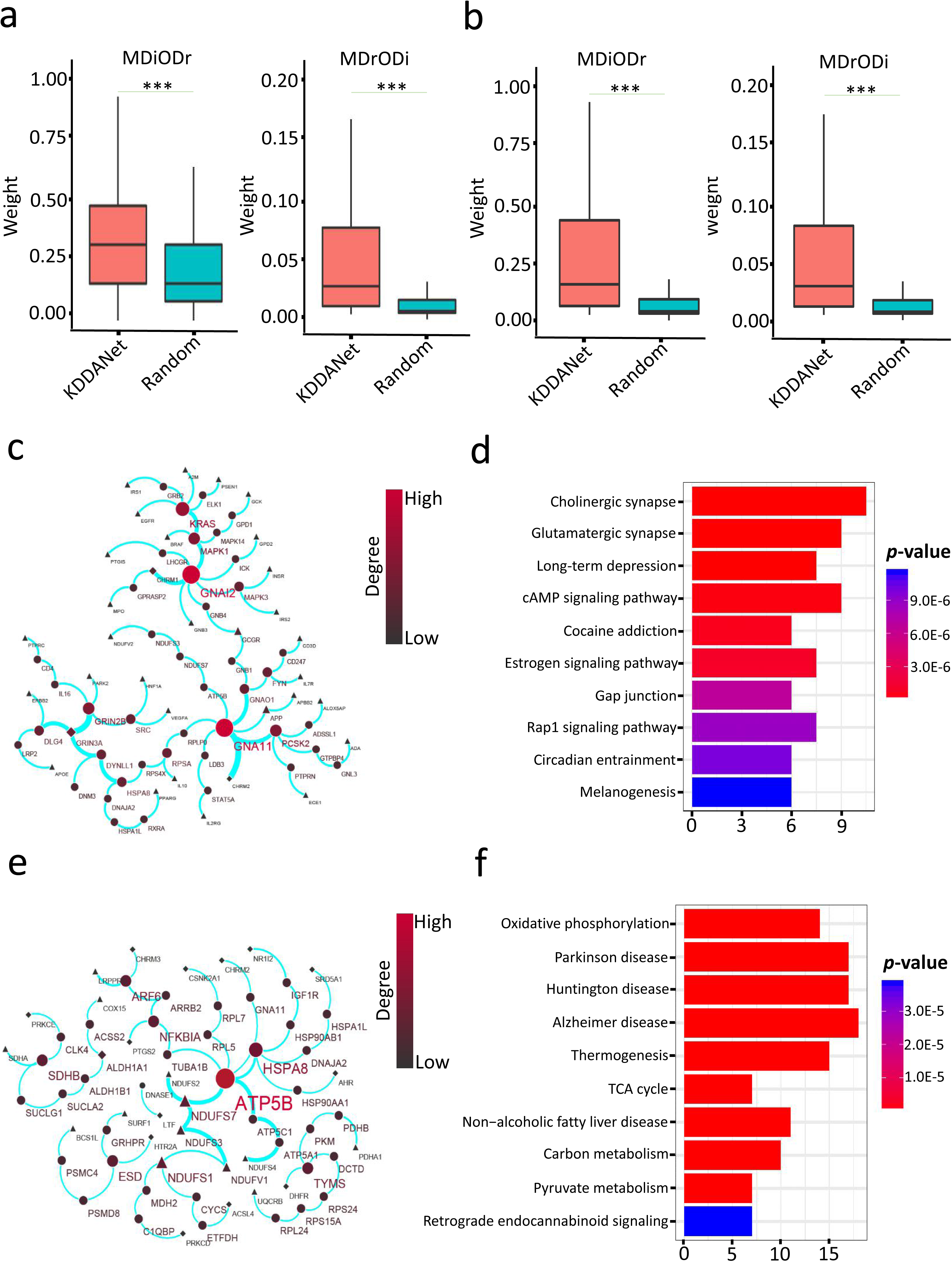
KDDANet uncovered the shared genes mediating multiple KDDAs. **a)** Boxplots demonstrating the distributions of weight values in KDDANet meta-subnetwork and random meta-subnetwork for MDiODr and MDrODi in SDrTDi context, *** *p*-value < 0.001, calculated by Wilcox signed rank test. **b)** Boxplots demonstrating the distributions of weight values of KDDANet meta-subnetwork and random meta-subnetwork for MDiODr and MDrODi in SDiTDr context, *** *p*-value < 0.001, calculated by Wilcox signed rank test. **c)** Shared meta-subnetwork mediating profenamine (DB00392)-neurological disease associations. **d)** Top 10 enriched KEGG terms of shared meta-subnetwork genes mediating profenamine-neurological diseases associations. **e)** Shared meta-subnetwork mediating the associations between GRACILE syndrome (603358) and 13 drugs. **f)** Top 10 enriched KEGG terms of shared meta-subnetwork genes mediating the associations between GRACILE syndrome and multiple drugs; In the meta-subnetworks, the size of a gene node was proportional to its network degree; the thickness of a network edge was proportional to its weight value.

We presented some examples for describing the capability of KDDANet in identifying shared genes mediating multiple KDDAs. For MDiODr, we selected two cases: I) profenamine (DB00392) and neurological disease associations and II) mirtazapine (DB00370) and cancer associations. As reported in CTD database, profenamine was associated with three neurological diseases, including Parkinsonian Disorders, Multiple Sclerosis and Alzheimer Disease. The shared meta-subnetwork contained 75 edges linking 43 unknown genes with three profenamine’s target genes and 30 neurological disease-related genes (Fig. 5c). The top 10 enriched KEGG terms were demonstrated in Fig. 5d. A majority of enriched KEGG terms, such as Cholinergic synapse and Glutamatergic synapses, have been reported dysfunction in neurological disorders and diseases ^83, 84^. Interestingly, we found that GNAI2 and GNA11 were two mostly shared genes linking profenamine with neurological diseases. These two genes were recently discovered involving in the pathological pathways of neurological diseases ^85, 86^. Mirtazapine was associated with 9 types of cancers, including Colorectal Neoplasms, Breast Neoplasms, Neuroblastoma, Glioma, Urinary Bladder Neoplasms, Stomach Neoplasms, Esophageal Neoplasms, Lung Neoplasms and Prostatic Neoplasms. Supplementary Figure 5e showed the shared meta-subnetwork that included 97 edges connecting 21 mirtazapine’s target genes with 33 cancer-related genes and 54 unknown genes. As expected, KEGG enrichment demonstrated that these genes were involved in cancer-related signaling pathways and played important roles in various cancers (Supplementary Figure 5f). Intriguingly, DRD4 and GRB2 were two mostly shared genes mediating the associations between mirtazapine and cancers (Supplementary Figure 5e). These two genes were involved in the oncogenesis of multiple types of cancers ^87, 88^.

For MDrODi, some interesting cases were also observed. For example, GRACILE syndrome, a metabolic disease, was associated with 13 drugs as recorded in CTD database. A shared meta-subnetwork including 66 genes and 61 edges were obtained for this disease (Fig. 5e). This meta-subnetwork contained 14 drug target genes and 13 GRACILE syndrome-related genes. The mostly shared gene was ATP5B, and the mostly significant enriched KEGG term was Oxidative phosphorylation (Fig. 5f). This was expected as GRACILE syndrome was a fatal inherited disorder caused by a mutation in an oxidative phosphorylation related gene, BCS1L^89^. It was also not surprising that the neurological diseases related genes were also enriched in this meta-subnetwork as patients with GRACILE syndrome had severe neurological problems^89^. Another example was the Keratoconus (148300), an ophthamological disease, which was associated with three different drugs, including acetaminophen, valproic acid and theophylline. Keratoconus and these three drugs shared a meta-subnetwork consisting of 54 genes and 51 edges (Supplementary Figure 5g). This meta-subnetwork included 7 Keratoconus-related genes, 11 drug target genes and 36 unknown genes. It was expected that HDAC2 had high weights with its partner genes in this meta-subnetwork as it was involved in notch signaling pathway which is downregulated in keratoconus ^90^. Consistent with the associations between collagen genes and keratoconus ^91^, a novel collagens coding gene, COL1A1, was captured in this meta-subnetwork as a highly shared gene. The enriched KEGG terms of this meta-subnetwork included Hippo signaling pathway (Supplementary Figure 5h) which has been reported involving in keratoconus corneas ^92^. Collectively, these results indicated that KDDANet can discover the shared genes mediating multiple KDDAs. The highly shared unknown genes can serve as potential candidate targets for drug repurposing.

## Discussion

To facilitate drug repurposing, various computational tools have been developed to uncover novel drug-disease associations ^93^. However, the potential genes mediating KDDA have been still not fully explored. Unveiling the hidden genes (missed from experiments) mediating KDDA become a great challenge for guiding novel target discovery and drug repurposing. In this work, we developed a novel computational tool, KDDANet, which integrated minimum cost network flow optimization, depth first searching and graph clustering algorithm to reveal hidden genes and modules mediating KDDA. KDDANet allowed for a global and systematic exploration of the hidden genes mediating KDDA. We applied KDDANet to unravel the subnetworks of genes mediating 53124 KDDAs. The comprehensive and system-level evaluations fully demonstrated the effective prediction capability and general applicability of KDDANet. Case studies on both AD and obesity showed that the subnetworks of genes identified by KDDANet were reliable and useful. Further validated by integrating analysis of genomic, transcriptomic, pharmacogenomic and survival data on primary tumors and cancer cell lines highlighted that KDDANet captured novel candidates from interactome mediating the associations between drugs and different types of cancer. Based on these, we concluded that the inferred subnetworks mediating KDDA can serve as genome-wide molecular landscapes for guiding drug repurposing and disease treatment. Insights learned from our predictions would also enable to help researchers to design repurposing drugs to reverse disease phenotypes via targeting key genes in the subnetwork mediating KDDA. An important capability of KDDANet method was that it can reveal the shared genes mediating multiple KDDAs. This provided more valuable guides for drug repositioning and disease treatment since the shared genes mediating multiple KDDAs were closely linked to the molecular basis of drug repurposing. We constructed a user-friendly online web tool (http://www.kddanet.cn), which allowed users to explore the subnetwork of genes mediating individual KDDA and the meta-subnetworks mediating multiple KDDAs (See Supplementary Note 7 as well as the Help section of our website for detailed description of the utility of an online web version of KDDANet). In summary, we presented a novel computational tool KDDANet and an online web source to decode the hidden genes mediating KDDA that had broad utility and application value in biomedical studies.

The hidden genes mediating KDDA predicted by KDDANet highlighted the power of integrative approaches to illuminate underexplored molecular processes mediating KDDA. In the future, the application value of our KDDANet tool in drug repurposing can be further improved from three aspects as follow: Firstly, the gene interactome used in KDDANet did not contain enough interactions between genome elements. In further studies, we would integrate other non-coding genome elements, especially long non-coding RNA and microRNA for constructing a comprehensive interactome. Secondly, integrating the biological networks from other omics layers, such as epigenomics, might have also further enhance the accuracy of KDDANet in discovering subnetworks and key genes mediating KDDA, and help us better to understand KDDA at multi-omics levels. The intrinsically capability of KDDANet to analyze large-scale heterogeneous interactome data containing tens of thousands of nodes and edges make it can well be suited to analyzing the accumulating data from ‘multi-omics’ technologies and biomedical research. Finally, KDDANet did not carry out predictions for KDDA when a drug’s target genes were unknown or when a disease has not been related to any known gene. For this, we planned to calculate the similarity scores between drugs and the similarity scores between diseases, and then integrate drugs without any target gene and diseases without any related gene to our flow network model by similarity scores.

## Methods

### Datasets

Five different types of gene networks, including HumanNet, HINT+HI2012, iRefIndex, MultiNet and STRINGv10, were used in our current study ^94, 95^, in which the nodes were represented by gene ID and connected by bidirectional edges (Supplementary Data 1). Drugs and their target genes were obtained from DrugBank 5.0.9 database (http://www.drugbank.ca/). In this study, we selected 4861 drugs with at least one known target that was contained in the gene networks for further analysis. In total, 2196 Known Drug Target Genes (KDTGs) included in the gene networks were connected to these drugs by 12014 interactions (Supplementary Data 2). Known Disease Related Genes (KDRGs) and classification of diseases were obtained by manually collecting Human Disease Network (HDN) from the previous study of Human Disease Network (HDN) ^33^ and DisGeNET v5.0 database ^96^. We focused on 1441 diseases with at least one related gene which was included in the gene networks for our study. In total, 16712 associations link these diseases to 1521 genes which were exist in the gene networks (Supplementary Data 3). The KDDAs were extracted from Comparative Toxicogenomics Database ^12^ (Supplementary Data 4). In this study, 53124 KDDAs were analyzed in which the drug had at least one target gene and the disease had at least one related gene contained in HumanNet. For simplicity and consistency, we converted different types of drugs, diseases, and gene nomenclatures to DrugBank drug ID, OMIM disease ID and NCBI Entrez gene ID for subsequent modeling and analysis.

The primary RNA-seq datasets of Alzheimer’s disease (AD) patients, obesity patients and normal individuals were downloaded from NCBI GEO Datasets under accession number of GSE53697, GSE81965 and GSE63887. After the SRA files were gathered, the archives were extracted and saved in FASTQ format using the SRA Toolkit. RNA-seq reads were trimmed using Trimmomatic software (http://www.usadellab.org/cms/?page=trimmomatic) with the following parameters “ILLUMINACLIP:TruSeq3-PE.fa:2:30:10 LEADING:3 TRAILING:3 SLIDINGWINDOW:4:15 MINLEN:36” (Version 0.36), and were further quality-filtered using FASTX Toolkit’s fastq_quality_trimmer command (http://hannonlab.cshl.edu/fastx_toolkit/) (Version 0.0.13) with the minimum quality score 20 and minimum percent of 80% bases that had a quality score larger than this cutoff value. The high-quality reads were mapped to the hg38 genome by HISAT2, a fast and sensitive spliced alignment program for mapping RNA-seq reads, with -dta paramenter (http://daehwankimlab.github.io/hisat2/). PCR duplicate reads were removed using Picard tools (http://broadinstitute.github.io/picard/) and only uniquely mapped reads were kept for further analysis. The expression levels of genes were calculated by StringTie (http://ccb.jhu.edu/software/stringtie/) (Version v1.3.4d) with -e -B -G parameters using Release 29 (GRCh38.p12) gene annotations downloaded from GENCODE data portal (https://www.gencodegenes.org/). To obtain comparable expression abundance estimation for each gene, reads mapped to hg38 were counted as FPKM (Fragments Per Kilobase Of Exon Per Million Fragments Mapped) based on their genome locations. Differential expression analysis of genes was performed by DESeq2 using the reads count matrix produced from a python script “prepDE.py” provided in StringTie website (http://ccb.jhu.edu/software/stringtie/).

TCGA cancer genomics datasets were directly downloaded via the UCSC Xena project data portal (https://xenabrowser.net/datapages/). As the DNA methylation datasets were quantified as beta value in the DNA probe level, we mapped Illumina Human Methylation 450 probe ID to gene name using HumanMethylation450 annotation file. If a gene was mapped by multiple probes, we considered the averaged signals of these probes as the methylation level of this gene. We used Wilcox signed rank test for differential analysis of gene expression and DNA methylation of KDDANet resulting subnetwork genes mediating the associations between drug and cancer. For this analysis, we only used the tumor samples which have adjacent normal tissue samples as control. Genomics of Drug Sensitivity in Cancer 1000 (GDSC1000) cancer cell line pharmacogenomic datasets were downloaded from GDSC website (https://www.cancerrxgene.org/gdsc1000/GDSC1000_WebResources/Home.html). Since this website provided only the CpG island methylation data, we downloaded the beta value matrix of probe-level DNA methylation from NCBI GEO Dataset under accession number of GSE68379 and then converted it to gene-level beta value matrix using the same method as TCGA DNA methylation data. Cancer Cell Line Encyclopedia (CCLE) pharmacogenomic datasets were directly downloaded from Broad Institute data portal (https://portals.broadinstitute.org/ccle/data).

### Construction of a unified flow network model

For each query drug and all its related diseases, the unified flow network model in SDrTDi context was built by integrating the query drug and all its related diseases into the gene network based on known drug-target relationships and gene-disease associations. As shown in Fig. 1, for the given query drug, we used it as source node (*S*) and integrated it into gene network by introducing the directed edges from it point to its target genes. For each its related disease, we mapped the disease to gene network by introducing the directed edges from its related genes point to it. We incorporated a sink node *T* and introduced a directed edge pointing from each disease to it. With these definitions, a unified flow network model was constructed as a complex heterogeneous graph *G* = (*V*, *E*), where *V* was the set of vertices and *E* was the set of edges. This graph included two types of edges (bidirectional and directed) and three types of nodes (drugs, genes, and diseases). Each edge was assigned with a weight and a capacity. Flow goes from a source node to a sink node through the graph edges. The assigning scheme of weight and capacity was illustrated as follow:

### Weight and capacity assigning scheme for network edges

*Edges between gene nodes.* Edges between gene nodes were weighted (*W_ij_*) to reflect the probability that two genes *g_i_* and *g_j_* were functionally linked in the biological processes. The weight value between *g_i_* and *g_j_* was derived from a Bayesian statistics approach by integrating diverse functional genomics datasets ^94^. Briefly, each dataset was benchmarked for its real capability of reconstructing known cellular pathways by measuring the likelihood that pairs of genes (linkages) were functionally connected conditioned on the experimental evidence, calculated as a log likelihood score *LLS* ^97^:

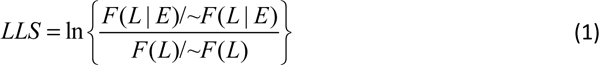

Where *F(L|E)* and *∼F(L|E)* represented the observed numbers of linkages *L* appear in the given experiment *E* between functionally annotated human genes interacting within the same pathway and between different pathways, respectively, *F(L)* and ∼*F(L)* denoted the total observed numbers of linkages between all annotated human genes interacting within the same pathway and between different pathways, respectively. The weight value (*W*) of a given linkage between two genes was produced by combining LLS score of each dataset using the formula:

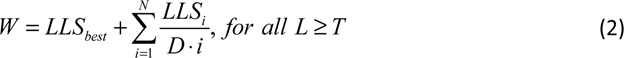

Where *LLS_best_* represented the highest *LLS* score for a linkage between two genes, *D* determined decay rate of the LLS score for additional evidence, and *i* was the order index of LLS scores for a given linkage between two genes, ranking starting from the second maximum LLS score with descending order of magnitude for all N remaining LLS scores. *T* represented a minimum threshold of LLS score to be considered. The values of *D* and *T* were empirically optimized to maximize overall performance on known GO annotations measured by AUPRC ^98^.

#### Edges between drug and drug’s target gene nodes

Edges between each drug and drug’s target gene nodes were weighted (*w_Si_*) to reflect the normalized reliability of the interaction between drug and target protein based on experimental and computational evidence. The weighting scheme was based on the predicted score of drug and target protein interaction ^25^. The weight value (*W_Si_*) was calculated as:

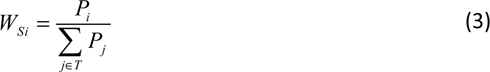

Where *T* denoted the set of each drug’s targets, *P_i_* denotes the predicted score between drug *S* and target *i*, *P_j_* denoted the predicted score between drug *S* and target *j*.

#### Edges between disease-related gene and disease nodes

Edges between disease-related gene and disease nodes were weighted (*w_jd_*) to reflect the normalized reliability of the linkage between disease and gene based on experimental and computational evidence. The weighting scheme was based on the predicted score of gene-disease association derived from MAXIF algorithm ^99^. We calculated the weight value (*W_jd_*) as follow:

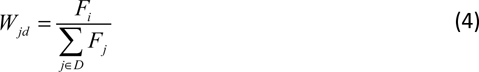

Where *D* denoted the set of genes linked to disease *d*, *F_i_* denoted the predicted score between gene *i* and disease *d*, *P_j_* denoted the predicted score between gene *j* and disease *d*.

#### Edges between disease and sink nodes

For each edge linking each disease *d* to sink node *T*, we assigned it a same weight value *W_dT_* = *1/N*, where *N* denoted the number of diseases linking to sink node.

We further defined for each edge in this heterogeneous network a capacity value that limited the flow quantity. For each edge connecting the query drug *S* to its target gene *i*, we assigned it a capacity *C_Si_* equal to *W_Si_*. For each edge linking the disease-related gene *j* to disease *d*, we assigned it a capacity *C_jd_* equal to *W_jd._* For each edge linking the disease *d* to sink node *T*, we assigned it a capacity *C_dT_* equal to *W_dT._* For other edges, we assigned them a capacity *C_ij_* = 1.

### Minimum cost flow optimization algorithm

With the purpose of identifying hidden genes mediating KDDA, we search for an possible solution that would (I) capture the subset of the query drug’s target genes which closely modulate all its related diseases by the disease-related genes without restrict to the prior KDDA genes, (II) determine hidden genes that were likely to be part of cellular pathways connecting the query drug’s target genes to all its related diseases but escaped detection by experiments, (III) give high priority to genes that lie on paths with highest probability connecting the query drug to all its related diseases without making constraints on the network structure. The rationality for proposing this solution including two aspects: I) this needed less computational time than that finding the highest probability subnetwork connects a query drug to each its related diseases at a time; II) this can effectively find the shared genes mediating multiple KDDAs. Inspired from the fact of water always flowing through a path of least resistance, we formulated this goal as a minimum cost flow optimization problem ^44, 100, 101^. Cost was defined as the negative log of the probability of an edge. Hence, minimizing the cost gave preference to highest-probability paths. Given the unified flow network, this problem can be expressed as a linear programming formula that minimized the total cost of the flow network while diffusing the most flow from query drug node to hypothetical sink node. Let *W_ij_*, *F_ij_* and *C_ij_* referred to the weight, flow, and capacity from node *i* to node *j*, respectively. The linear programming formula can be written as follow.

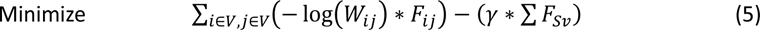

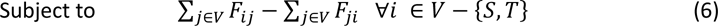

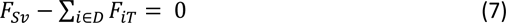

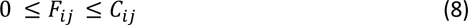

Where *V* denoted a set of nodes included in the flow network; *S* denoted the query drug and *v* denoted its target gene. The parameter gamma (*y*) controlled the size and the quality of the optimized subnetwork. The first component of this formula, *∑_i∈V,j∈V_*(− log(*W_ij_*) ∗ *F_ij_)*, ensured minimizing the network cost that given priority to obtain highest probability gene subnetwork, at the same time, the second component, *−(y ∗ ∑ F_Sv_)*, indicated maximizing the total flow across entire network. This optimization problem can be efficiently solved using primal simplex method provided in Mixed Integer Linear Programming (MILP) solver (http://lpsolve.sourceforge.net/). The solution *argmin_fij > 0_ ∑_i∈V,j∈V_*(− log(*w_ij_*) ∗ *f_ij_*) − (*y ∗ ∑ f_Sv_*) obtained the highest probability subnetwork mediating the associations between query drug and all its related diseases.

### Depth first searching and Markov clustering (MCL)

Once the highest probability subnetwork mediating the query drug (disease) with all its related diseases (drugs) was obtained, we implemented depth first searching ^102^ on this subgraph to find the subnetwork made up of all paths linking the query drug (disease) to each it’s related disease (drug). All genes in the solution were ranked by the amount of flow they carry. The more flow that passes through a protein, the more important it was in mediating KDDA. After obtaining the subnetwork mediating individual KDDA, Markov clustering (MCL) (https://micans.org/mcl/) was employed to further discover gene modules mediating KDDA by using the flow quantities through edges of subnetwork as weight values.

### Application KDDANet to SDiTDr context

To apply KDDANet in SDiTDr context, the query disease, and a set of all its related drugs were mapped into gene network by disease-related genes and drug target genes. For constructing flow network model, the weights and capacities of network edges can be assigned using the similar method as described in SDrTDi context by substituting query drug with query disease and using query disease as source node (*S*), substituting query drug related diseases with query disease related drugs, substituting query drug’s target genes with query disease related genes, substituting disease related genes with drug’s target genes, and incorporating a sink node *T* and introducing a directed edge pointing from each drug to it. Implementing minimum cost flow optimization, depth first searching and Markov clustering (MCL) on the unified flow network was same as SDrTDi context.

### Performance evaluation

As the true genes mediating KDDA are poorly understood, there was no perfect way to assess the prediction results. The predictive performance of KDDANet was evaluated as follow: I) Based on the hypothesis that the larger functional similarity between a gene and known KDDA genes, the higher probability this gene was positive one mediating KDDA, we compiled a standard set of positive and negative KDDA genes for each KDDA resulting subnetwork to unbiasedly evaluate the performance of KDDANet using the following strategy:

I) For each gene in the KDDA resulting subnetwork, we first calculated the mean functional similarity scores of it with KDTGs and KDRGs by our previous published method using Gene Ontology (GO), KEGG pathways and InterPro annotation as functional terms ^103^.

II) We performed a permutation test by randomly producing KDTGs and KDRGs 1000 times to compute the empirical significance level of functional similarity. We selected the genes having significant functional similarities with both KDTGs and KDRGs from KDDA resulting subnetwork as positive KDDA genes using the criterion that the similarity score was larger than 95th percentiles of the simulated background distributions. The other genes were considered as negative KDDA genes.

Based on this gold standard, Receiver Operating Characteristic (ROC) and Precision-Recall (PR) curves were produced and the areas of under curve of ROC and PR (AUROC and AUPRC) were calculated for gene list ranked by flow amount of each KDDANet resulting subnetwork. For comparison with PPA, SNPLS, comCHIPER and DGPsubNet, we used the default parameters as mentioned in their published papers. We considered that all genes contained in a co-module mediating KDDA of each drug and disease pair in the co-module. The genes were ranked by probability score mediating a KDDA. For PPA and SNPLS, the probability score of a gene *g* mediating the association between drug *i* and a type of cancer *j* was defined as averaged prediction score across all cell lines of this type of cancer ^40, 41^. For comCHIPER, the probability score of a gene *g* mediating the association between drug *i* and disease *j* was defined as the sum of the products of posterior indicator probabilities across all co-modules ^42^. For DGPsubNet, we defined the probability score of a gene *g* mediating the association between drug *i* and disease *j* as the calculated *z-score z_ij_* ^43^. The areas of under curve of ROC and PR (AUROC and AUPR) were calculated using R package of PRROC. KDDANet resulting subnetwork visualization was carried out using Cytoscape software (https://cytoscape.org/). KEGG pathway enrichment analysis was performed by clusterProfiler R package ^104^. The visualization of results was carried out in R software. Case studies were conducted in SDrTDi context unless specifically indicated.

## Data Availability

The authors declare that data supporting the findings of this study are available within the paper and its supplementary information files.

## Code availability

The code of KDDANet was freely available at https://github.com/huayu1111/KDDANet.

## Acknowledgements

H.Y. was supported by Project funded by China Postdoctoral Science Foundation (Grant No. 2018M642441), Zhejiang Natural Science Foundation Projects of China (Grant No. LQ21C120002) and High-Performance Computing Platform in Center of Cryo-Electron Microscopy of Zhejiang University.

## Author Contributions

H.Y. and L.L. conceived the original research plans, analyzed the data, developed software, and wrote the article; L.L and H.Y developed the web server. H.Y., supervised the experiments; H.Y. and L.L. contributed equally to this work. J.Z., C.L., and M.C. provided helpful suggestions and computational support. All authors revised and confirmed the paper.

