## Supplementary Note; Supplementary Figure; Supplementary Data for "Genome-wide discovery of hidden genes mediating known drug-disease association using KDDANet"

[1] Center for Stem Cell and Regenerative Medicine, Department of Basic Medical Sciences, and The First Affiliated Hospital, Zhejiang University School of Medicine, Hangzhou, Zhejiang, China. Institute of Hematology, Zhejiang University, Hangzhou, Zhejiang, China.

[2] Department of Human Genetics, and Women's Hospital, Zhejiang University School of Medicine, Hangzhou, China

[3] College of Life Sciences, Zhejiang University, Hangzhou, China.

[4] Lead contact

[#] These authors contributed equally.

#### Supplementary Figure Legends

**Supplementary Figure 1:** Performance evaluation of KDDANet method.

**Supplementary Figure 2:** General applicability of KDDANet method.

**Supplementary Figure 3:** Mechanistic relevance of KDDANet prediction results.

**Supplementary Figure 4:** KDDANet provided novel molecular insights on KDDAs related to cancer.

**Supplementary Figure 5:** KDDANet uncovered the shared genes mediating multiple KDDAs.

#### Supplementary Note Legends

**Supplementary Note 1:** A mini review of state-of-the-art computational tools for facilitating drug repurposing.

**Supplementary Note 2:** Enrichment analysis of hidden genes mediating KDDA in the KDDANet resulting subnetworks.

**Supplementary Note 3:** Method for enrichment analysis.

**Supplementary Note 4:** Method for selecting a suitable $\gamma$ value.

**Supplementary Note 5:** KEGG pathway enrichment of KDDANet resulting subnetworks.

**Supplementary Note 6:** Analysis of KDDANet resulting subnetworks mediating the associations between drugs and cancer using cancer-omics' datasets.

**Supplementary Note 7:** An online web server for KDDA decoding.

#### Supplementary Data Legends

**Supplementary Data 1:** Five different types of gene interaction networks, including HumanNet, HINT+HI2012, iRefIndex, MultiNet and STRINGv10 used in this study.

**Supplementary Data 2:** Drugs and their target genes used in this study.

**Supplementary Data 3:** Known diseases and their related genes used in this study.

**Supplementary Data 4:** 53124 KDDAs used in this study.

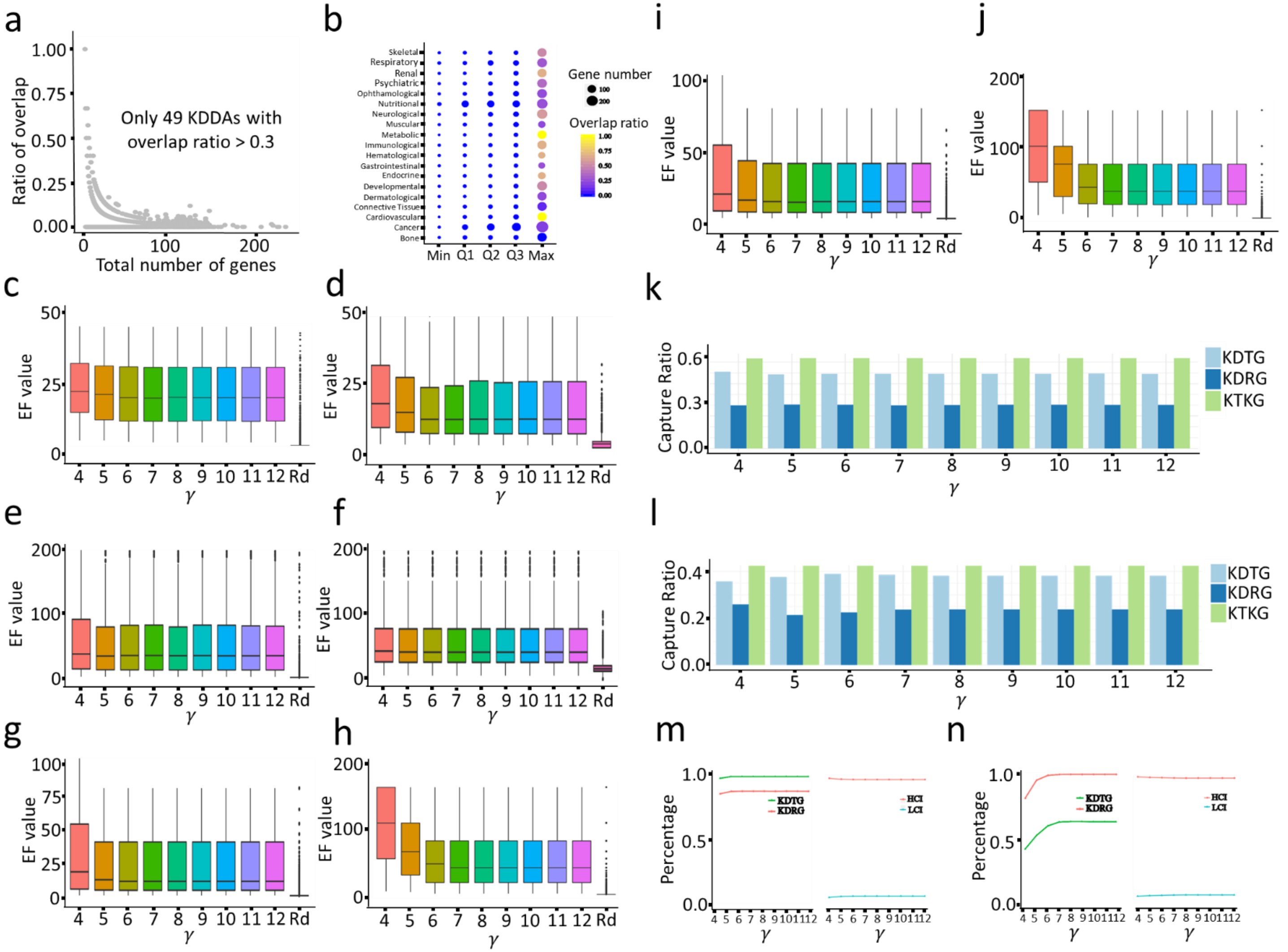

a
b
i
j
1.00
0.75
0.50
0.25
0.00
Ratio of overlap
Only 49 KDDAs with
overlap ratio > 0.3
0
100
200
Total number of genes
Skeletal
Respiratory
Renal
Psychiatric
Ophthamological
Nutritional
Neurological
Muscular
Metabolic
Immunological
Hematological
Gastrointestinal
Endocrine
Developmental
Dermatological
Connective Tissue
Cardiovascular
Cancer
Bone
Min
Q1
Q2
Q3
Max
Gene number
Overlap ratio
EF value
γ
4 5 6 7 8 9 10 11 12 Rd
c
d
e
f
g
h
k
l
m
n
Capture Ratio
KDTG
KDRG
KTKG
Percentage
HCI
LCI

**Supplementary Figure 1: Performance evaluation of KDDANet method. a)** Scatter plot demonstrating the distribution of overlap ratio between KDTGs and KDRGs. The x-axis denoted the total number of KDTGs and KDRGs mediating a KDDA. **b)** Scatter plot demonstrating the Minimum (Min), 25th Quantile (Q1), 50th Quantile (Q2), 75th Quantile (Q3) and Maximum (Max) of overlap ratio and gene number distribution of 53124 KDDAs in different types of diseases; the size and color of point denoted gene number and overlap ratio, respectively. **c)** Boxplot demonstrating the enrichments of KDTGs in KDDANet resulting subnetwork with different $\gamma$ settings in SDrTDi context, using random permutation as control (Rd). **d)** Similar to as **c)**, demonstrating the enrichments of KDRGs. **e)** Similar to **c)**, demonstrating the enrichments of KDTGs in SDiTDr context. **f)** Similar to **d)**, demonstrating the enrichments of KDRGs in SDiTDr context**. g)** Boxplot demonstrating the enrichment of KTKGs in SDrTDi context, using random permutation as control (Rd). **h)** Similar to **g)**, demonstrating the enrichment of KTKGs in SDiTDr context. **i)** Similar to **g)**, demonstrating the enrichment of NTKGs. **j)** Similar to **h)**, demonstrating the enrichment of NTKGs. **k)** The KDDANet capture ratio of KDTGs, KDRG and KTKGs with different γ setting in SDrTDi context. **l)** The KDDANet capture ratio of KDTGs, KDRGs and KTKGs with different γ setting in SDiTDr context. **m)** Line chart demonstrating the percentage of KDTGs and KDRGs that were incorporated into the KDDANet resulting subnetwork, as well as the percentage of low probability edges with weight smaller than 0.3 in SDrTDi context (HCI, high confidence interaction; LCI, low confidence interaction). **n)** Similar to **m)**, demonstrating the percentages in SDiTDr context. In panels **c**-**j**, the centre of the box plots represents the median value and the lower and upper lines represent the 25% and 75% quantile, respectively. The whiskers stand for 1.5 × interquartile range below the lower quartile and above the upper quartile. The points located outside of whiskers denote outliers.

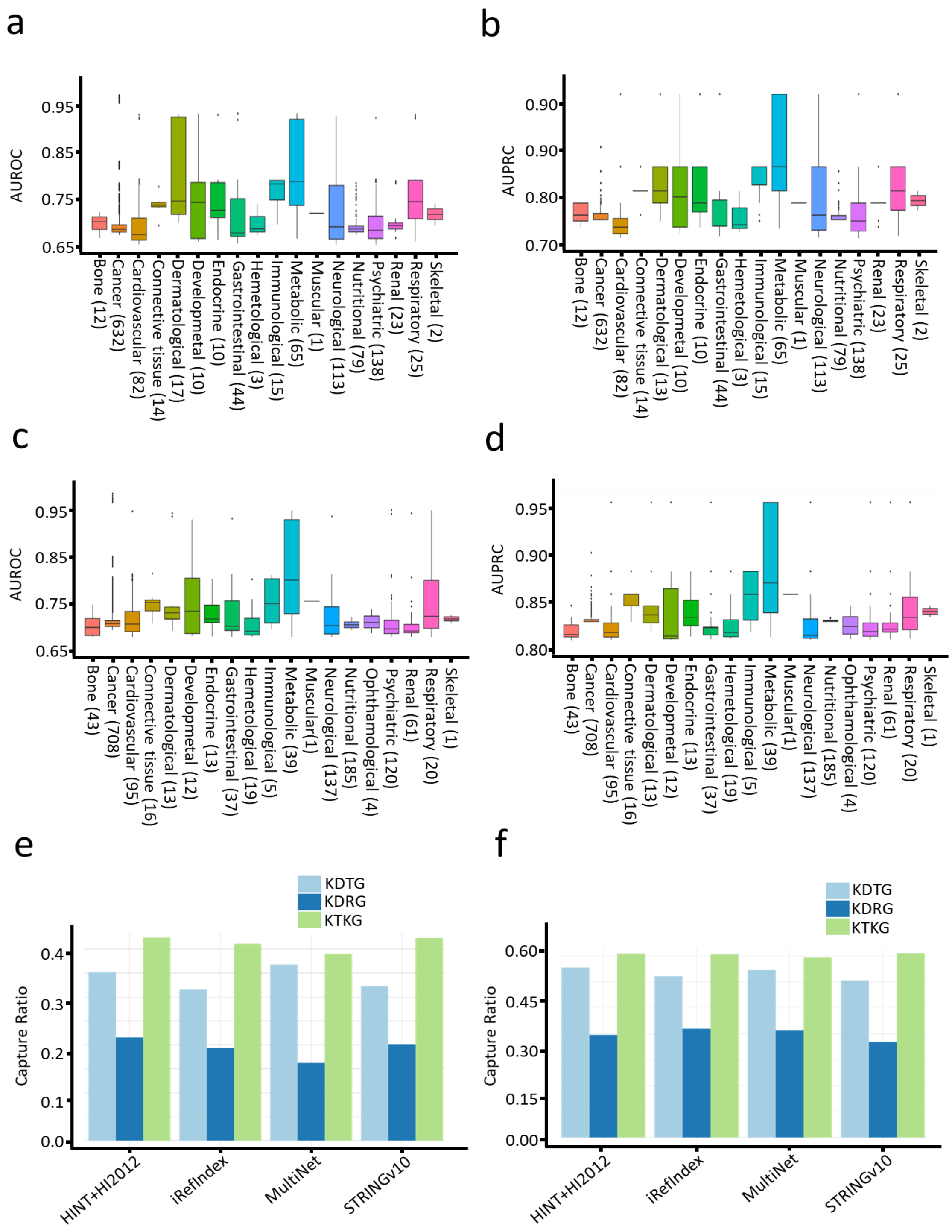


**Supplementary Figure 2: General applicability of KDDANet method. a)** and **b)** AUROC and AUPRC of KDDANet across different types of diseases in SDrTDi context. **c)** and **d)** Similar to **a)** and **b)** in SDiTDr context; Values in parentheses denotes the number of KDDAs. **e)** The KDDANet capture ratio of KDTGs, KDRGs and KTKGs using different types of networks in SDiTDr context. **f)** Similar to **e)** in SDrTDi context. In panels **a**-**d**, the centre of the box plots represents the median value and the lower and upper lines represent the 25% and 75% quantile, respectively. The whiskers stand for 1.5 × interquartile range below the lower quartile and above the upper quartile. The points located outside of whiskers denote outliers.

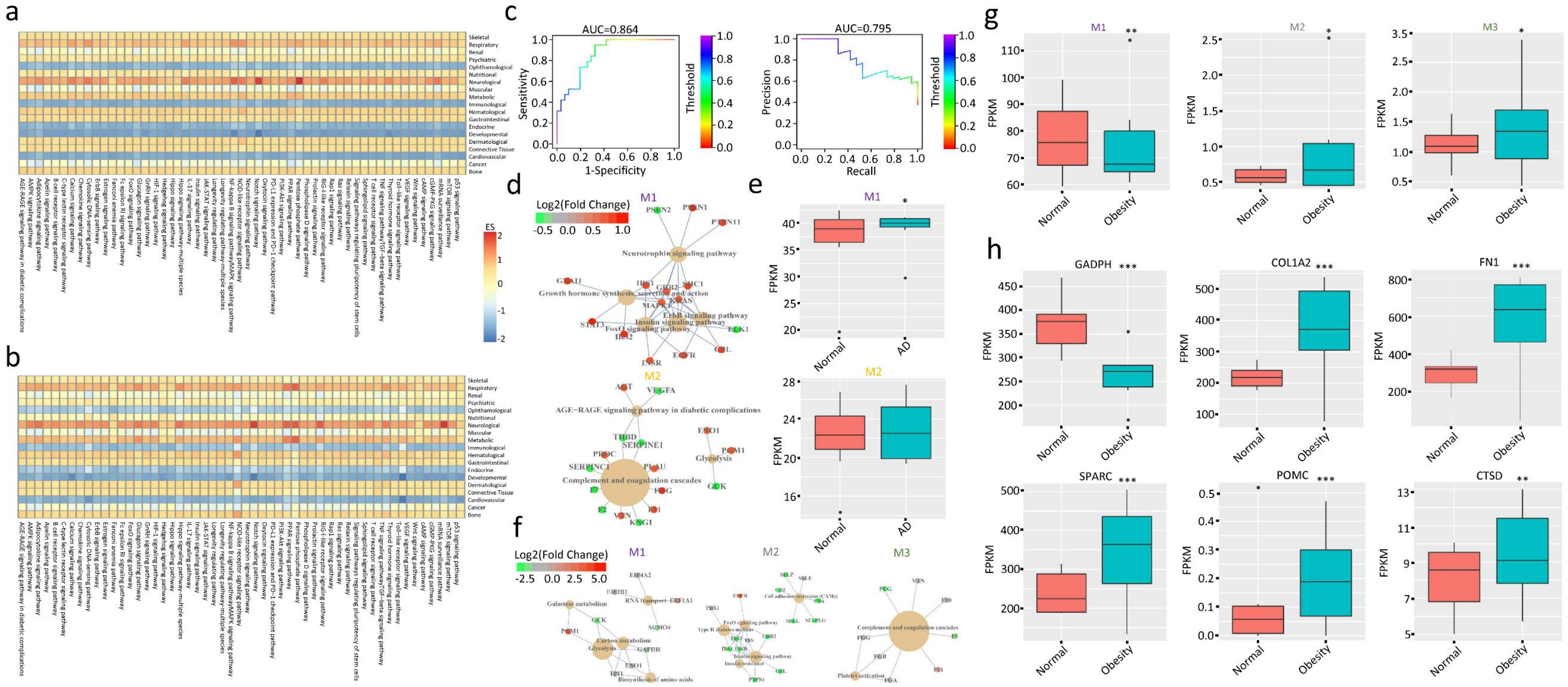


**Supplementary Figure 3: Mechanistic relevance of KDDANet prediction results**. **a)** Enrichment Score (ES) heatmap of 53 KEGG pathways on 19 different types of diseases in SDrTDi context. **b)** Similar to **a)**, in SDiTDr context. A two-step procedure was used to calculate ES. First, for each disease type, we calculated the frequency of KDDA resulting subnetworks enriched on 53 KEGG pathways and obtained a frequency matrix. Then, for each pathway, we normalized the frequency matrix to a *z*-score matrix. The value in *z*-score matrix was defined as ES. **c)** ROC and PR curves of KDDANet resulting subnetwork mediating phylloquinone-Alzheimer's disease (AD) association. **d)** Pathway-gene relationship network demonstrating the enriched KEGG terms and their related genes for module M1 and M2 mediating phylloquinone-AD association. **e)** Averaged expression level of module M1 and M2 genes in normal individuals and AD patients, * $p$-value < 0.05, calculated by Mann-Whitney U test. **f)** Similar to **d)** pathway-gene relationship network demonstrating the enriched KEGG terms and their related genes for module M1, M2 and M3 mediating heparin-obesity association. **g)** Averaged expression level of module M1, M2 and M3 genes in normal individuals and obesity patients, ** $p$-value < 0.01, * $p$-value < 0.05, calculated by Mann-Whitney U test. **h)** Expression level of GADPH, COL1A2, FN1, SPARC, POMC and CTSD in normal individuals and obesity patients; *** $p$-value < 0.001, ** $p$-value < 0.01, calculated by Mann-Whitney U test. In the pathway-gene relationship networks, the size of a KEGG term node was proportional to its $p$-value in enrichment analysis; The color of a gene node denoted its fold change of expression level between normal individuals and patients. Fragments Per Kilobase Of Exon Per Million Fragments Mapped, FPKM. In panels **e**, **g and h**, the centre of the box plots represents the median value and the lower and upper lines represent the 25% and 75% quantile, respectively. The whiskers stand for 1.5 × interquartile range below the lower quartile and above the upper quartile. The points located outside of whiskers denote outliers.

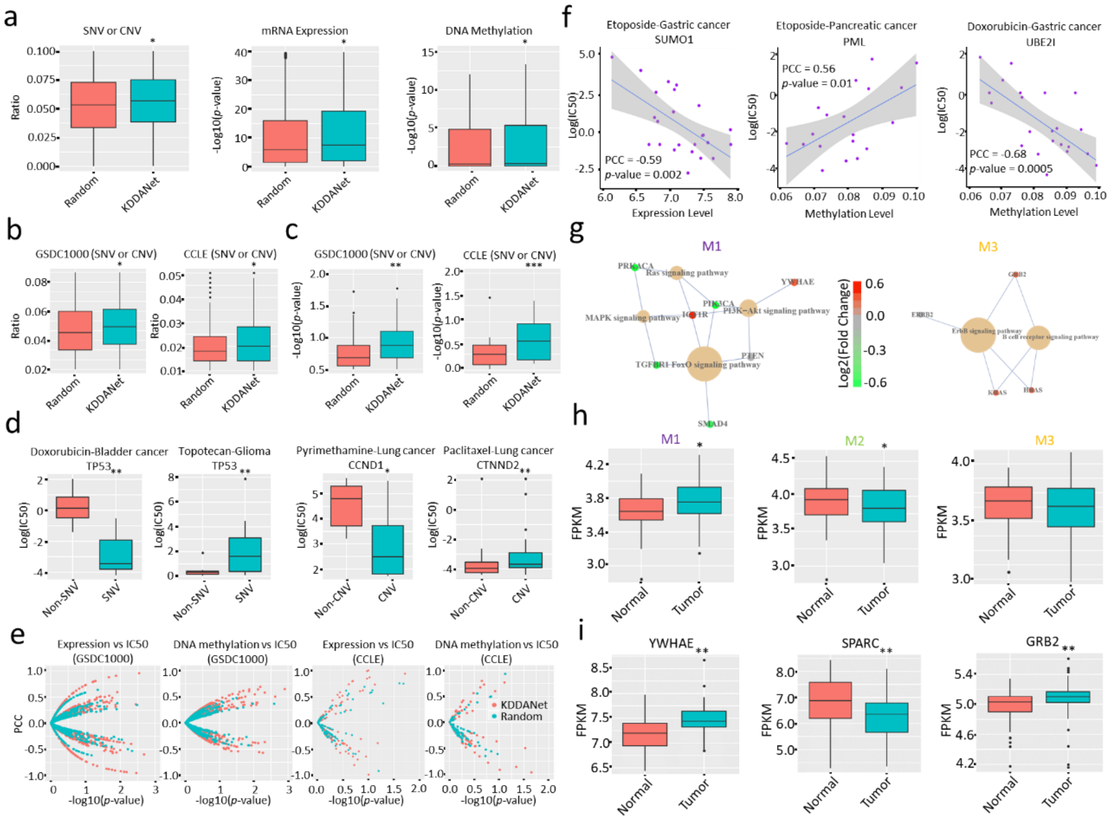


**Supplementary Figure 4: KDDANet provided novel molecular insights on KDDAs related to cancer. a)** The ratio of TCGA tumor samples harboring SNVs or CNVs of KDDANet resulting subnetwork genes mediating the associations between drugs and cancer and randomly selected genes, and the overall statistic difference of mRNA expression and DNA methylation of KDDANet resulting subnetwork genes mediating the associations between drugs and cancer (randomly selected genes) between tumor samples and normal samples, respectively; * *p*-value < 0.05, calculated by Mann-Whitney U test. **b)** The ratio of cancer cell lines harboring SNVs or CNVs of KDDANet resulting subnetwork genes mediating the associations between drugs and cancer and randomly selected genes, * *p*-value < 0.05, calculated by Mann-Whitney U test. **c)** Boxplots demonstrating the overall statistic difference between IC50 values of anti-cancer drugs on cell lines harboring SNVs or CNVs of KDDANet resulting subnetwork genes mediating the associations between drugs and cancer and cell line without SNVs and CNVs of these genes, *** *p*-value < 0.001, ** *p*-value < 0.01, calculated by Mann-Whitney U test. **d)** Representative examples demonstrating the difference between IC50 values of anti-cancer drugs on cell lines harboring SNVs or CNVs of KDDANet resulting subnetwork genes mediating the associations between drugs and cancer and cell lines without SNVs and CNVs of these genes, ** *p*-value < 0.01, * *p*-value < 0.05, calculated by Mann-Whitney U test. **e)** Scatter plot demonstrating the correlation between expression abundances/methylation levels of KDDANet resulting subnetwork genes mediating the associations between drugs and cancer (randomly selected genes) and half maximal inhibitory concentration (IC50) values of anti-cancer drugs in both GSDC1000 and CCLE cancer cell lines, respectively; PPC, Pearson Correlation Coefficients; *p*-value was calculated by cor.test function in R software. **f)** Representative examples demonstrating the correlation between the expression abundances/methylation levels of KDDANet resulting subnetwork genes mediating the associations between drugs and cancer and IC50 values of anti-cancer drugs in both GSDC1000 and CCLE cancer cell lines, respectively. **g)** Pathway-gene relationship network demonstrating the enriched KEGG terms and their related genes for M1 and M3 mediating sotalol (DB00489)-prostate cancer (176807) association. **h)** Averaged expression level of module M1, M2 and M3 genes in prostate cancer tumor samples and adjacent normal tissue samples, * *p*-value < 0.05, calculated by Wilcox signed rank test. **i)** Expression level of YWHAE, SPARC and GRB2 in prostate cancer tumor samples and adjacent normal tissue samples, ** *p*-value < 0.01, calculated by Wilcox signed rank test. In panels **a**-**d**, **h** and **i**, the centre of the box plots represents the median value and the lower and upper lines represent the 25% and 75% quantile, respectively. The whiskers stand for 1.5 × interquartile range below the lower quartile and above the upper quartile. The points located outside of whiskers denotes outliers.

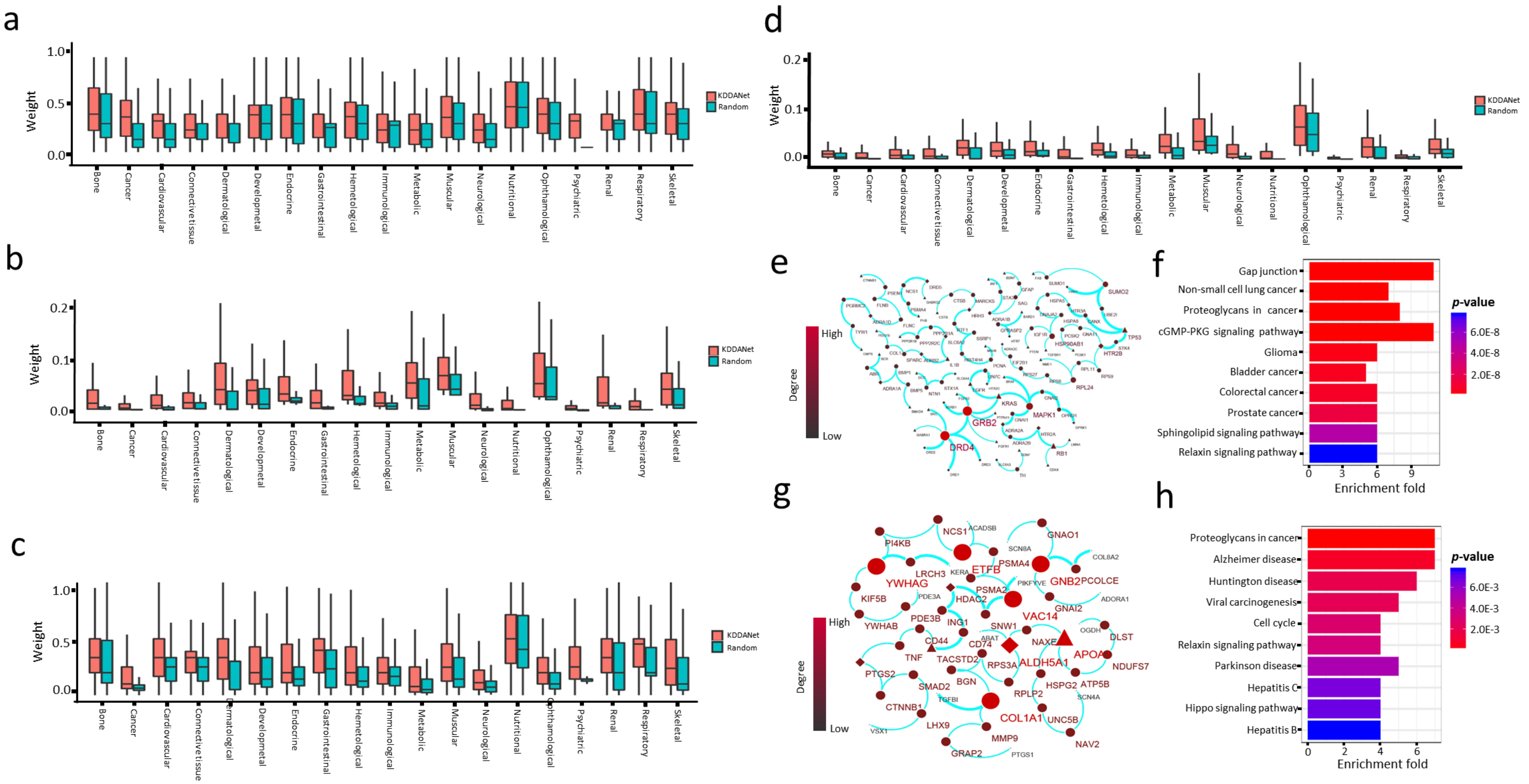


**Supplementary Figure 5: KDDANet uncovered the shared genes mediating multiple KDDAs. a)** Boxplots demonstrating the distributions of weight values in KDDANet meta-subnetwork and random meta-subnetwork for MDiODr in different types of diseases in SDrTDi context. **b)**

Boxplots demonstrating the distributions of weight values in KDDANet meta-subnetwork and random meta-subnetwork for MDrODi in different types of diseases in SDrTDi context. **c)** Similar to **a)** in SDiTDr context; **d)** Similar to **b)** in SDiTDr context. **e)** Shared meta-subnetwork mediating the associations between mirtazapine (DB00370) and 9 types of cancers. **f)** Top 10 enriched KEGG terms of shared genes of meta-subnetwork mediating the associations between mirtazapine (DB00370) and 9 types of cancers. **g)** Shared meta-subnetwork mediating the associations between Keratoconus (148300) and three drugs. **h)** Top 10 enriched KEGG terms of shared genes of meta-subnetwork mediating the associations between Keratoconus (148300) and three drugs. In panels **a**-**d**, the centre of the box plots represents the median value and the lower and upper lines represent the 25% and 75% quantile, respectively. The whiskers stand for 1.5 × interquartile range below the lower quartile and above the upper quartile. The points located outside of whiskers denotes outliers.

#### Supplementary Note 1: A mini review of state-of-the-art computational tools for facilitating drug repurposing.

With the rapid accumulation of biomedical data, various computational methods, including machine learning, similarity computation and network-based models, have been proposed for facilitating drug repurposing. We here provided a mini summary of state-of-the-art computational approaches designed for predicting drug-drug interactions, drug side-effects, drug combinations, drug indications, drug-target interactions, disease-related genes, disease modules and constructing disease-disease relationship network. For forecasting pharmacokinetic and pharmacodynamic drug-drug interactions and their associated recommendations, Gottlieb et al. designed "INDI", a similarity measure-based logistic regression classifier [1]. In addition, Huang et al. proposed a metric "S-score" that measures the strength of connection between drug targets in the protein-protein interaction network and integrates drug clinical side effects to predict pharmacodynamic drug-drug interactions [2]. For identifying drug side-effects, Atias et al. and Shaked et al. designed the network-based approaches [3,4]. Wang et al. developed a multilabel classifier by combining chemical structure and gene expression features to predict drug side-effects [5]. Tatonetti et al. presented an adaptive data-driven approach for predicting drug side-effects and drug interaction side-effects [6]. Duffy et al. used the tissue-specific genetic features for drug side-effects prediction in clinical trivials [7]. Kuhn et al. integrated phenotypic data obtained during clinical trials with known drug-target relations to identify overrepresented protein-side effect combinations [8]. For screening novel drug combinations, Zhao et al. explored various molecular and pharmacological features of drugs and showed that combinations of such features were an effective predictor [9]. In addition, the network-based methods have been proposed as a promising tool to identify novel drug combination [10,11]. Interestingly, Kuenzi et al. developed a deep-learning model of visible neural network, called DrugCell, to predict drug response and synergy in human cancer cells [12]. To predict novel drug-disease associations and identify drug indications, Pacini et al. developed a R/Cytoscape package, which utilized the correlation of gene expression profiles to infer whether a query drug can be repurposed for a disease [13]. Gottlieb et al. developed PREDICT, which scored a possible drug-disease association by combining multiple drug-drug and disease-disease similarity measures [14]. Guney et al. proposed an unsupervised and unbiased network-based proximity measure between drug targets and disease proteins for scoring drug-disease associations [15]. Cheng et al. further showed that the integration of network proximity-based approach and large-scale patient-level longitudinal data complemented by mechanistic in vitro studies offered an effective platform by which to identify and validate novel drug-disease associations and drug indications [16]. Besides, Gilvary and Elkhader developed a machine learning and network framework to discover new indications for small molecules [17]. For identifying novel drug targets, Paolini et al presented the global mapping of pharmacological space that can identify confidently the human drug targets [18]. Yamanishi et al. proposed a new statistical method that formalized the drug-target interaction inference as a supervised learning problem for a bipartite graph and integrated chemical and genomic spaces into a unified pharmacological space [19]. Campillos et al. used the side-effect similarity to identify novel drug targets [20]. Perlman et al. developed "SITAR", which incorporated multiple drug-drug and gene-gene similarity measures for scoring drug and gene associations [21]. Yu et al employed both support vector machine and random forest for identifying multiple drug and target interactions [22]. Luo et al. developed a computational pipeline, called DTINet, to predict novel drug–target interactions from a constructed heterogeneous network [23]. The network propagation algorithm has been also proposed for drug target identification [24]. Iskar et al. identified and characterized drug-induced transcriptional modules using microarray data [25]. Silberberg et al. further proposed a novel method to identify drug response pathways by combining drug targets, drug response expression profiles, and the human physical interaction network [26]. For discovering the disease-related genes and disease modules, multiple computational approaches have been designed [27-36]. For example, Wu et al. designed CIPHER for identifying human disease genes that integrated human protein-protein interactions, disease phenotype similarities, and known gene-phenotype associations to capture the complex relationships between phenotypes and genotypes [27]. Gottlieb et al. developed “PRINCIPLE”, which employed the classical network propagation algorithm for associating genes with diseases [30]. Ghiassian et al. designed “DIAMOnD” for detecting disease modules based on systematic analysis of connectivity patterns of disease proteins in the human interactome [31]. Particularly, Vinayagam et al. identified both disease genes and drug targets based on controllability analysis of the directed human protein interaction network [36]. Menche et al. used the biomedical literature database and incomplete interactome to construct disease-symptom and disease-disease relationship networks [37,38]. Moreover, Hofree et al. introduced network-based stratification (NBS) to uncover tumor subtypes by integrating somatic tumor genomes with gene networks [39]. Collectively, these computational methods helped our understanding of the pathology of diseases and mechanisms of drug actions, and thus promoted the process of drug repurposing.

#### Supplementary Note 2: Enrichment analysis of hidden genes mediating KDDA in the KDDANet resulting subnetworks.

We first tested whether the KDTGs and KDRGs are contained in KDDANet resulting subnetworks by performing enrichment analysis with a range of $\gamma$ setting from 4 to 12 with a step 1. For comparing with background, we carried out a permutation test by producing random subnetworks with the same number of genes. We found that the KDTGs and KDRGs have significant enrichment in the resulting subnetworks against random subnetworks (Supplementary Figure 1c-1f). This indicated that KDDANet can effectively capture KDTGs and KDRGs. To examine whether KDDANet can capture true genes mediating KDDA, we introduced two concepts: “known true KDDA genes (KTKGs)” and “novel true KDDA genes (NTKGs)” (see Main Text for definition). Based on these definitions, we then performed enrichment analysis to check whether the KTKGs have captured in our prediction results. As shown in Supplementary Figure 1g and Supplementary Figure 1h, the subnetworks outputted by KDDANet have obviously higher enrichment of KTKGs when compared with the permutation subnetworks. We further performed enrichment analysis to test whether KDDANet can uncover NTKGs. We found that the resulting subnetworks have still higher enrichment of NTKGs than random permutation subnetworks (Supplementary Figure 1i and Supplementary Figure 1j). We removed the link of each KTKG with drug and disease from the flow network to test the recoverability of KDDANet. Interestingly, we observed that the capture ratio of KTKG in KDDANet resulting subnetwork is higher than KDTG and KDRG (Supplementary Figure 1k and Supplementary Figure 1l). This suggested that KDDANet favor to capture more NTKGs than KDTGs and KDRGs.

#### Supplementary Note 3: Method for enrichment analysis.

We carried out fold enrichment analysis against full gene network for each predicted KDDA subnetwork to evaluate whether KDDANet can capture true genes mediating KDDA. The fold of enrichment was calculated by the following formula:

$$\frac{p}{q} \times \frac{M}{N} \tag{1}$$

where, *p* was the number of "KDTGs"/"KDRGs" or the number of true positive genes ("KTKGs"/"NTKGs") mediating a given KDDA; *q* was the number of inferred genes in the resulting KDDA subnetwork; *M* denoted the number of all "KDTGs"/"KDRGs" or the number of all true positive genes ("KTKGs"/"NTKGs") in the gold standards and *N* represented the number of all genes in full network.

#### Supplementary Note 4: Method for selecting a suitable $\gamma$ value.

As the size and quality of KDDANet output subnetwork depended on a parameter $\gamma$. Higher $\gamma$ values will identify more links between the source node and the sink node but with lower confidence. To obtain suitable values for $\gamma$, we run KDDANet with $\gamma$ values ranging between 4 and 12 with a step of 1. For each of the output subnetworks, we computed the percentage of input, namely KDTGs and KDRGs, that were incorporated into the network, as well as the percentage of low probability edges with weight smaller than 0.3. As shown in Supplementary Figure 1m and Supplementary Figure 1n, we observed that the percentages for KDTGs, KDRGs and low probability edges contained in resulting subnetworks became stable when γ = 6 and 8 for SDrTDi and SDiTDr context, respectively. We therefore selected γ = 6 for SDrTDi and γ = 8 for SDiTDr as they were the minimal values with which a significant fraction of the input was incorporated while the percentage of low probability edges remained small in respective contexts.

#### Supplementary Note 5: KEGG pathway enrichment of KDDANet resulting subnetworks.

We asked whether the enriched functions of KDDANet subnetworks were mechanistically related to KDDA by carrying out a global enrichment of all predicted KDDA subnetworks against 53 classical KEGG pathways. We observed that p53 signaling pathway had relatively higher enrichment in KDDA subnetwork linking drug with cancer when comparing with other pathways in both SDrTDi and SDiTDr contexts (Supplementary Figure 3a and Supplementary Figure 3b). For neurological disease, we found that the notch-signaling pathway had significant enrichment in SDrTDi and SDiTDr context (Supplementary Figure 3a and Supplementary Figure 3b). This was consistent with the previous discovery that showed the association between notch-related pathways and neurological disorders [40]. Interestingly, we found that ophthamological disease, immunological disease, endocrine disease, developmental disease, and cardiovascular diseases have consistent less enrichment on all KEGG pathways than other types of diseases (Supplementary Figure 3a and Supplementary Figure 3b). This indicated that the drugs developed to cure these types of diseases cannot be well explained by the known pathways and the uncovered molecular interactions mediating KDDAs for these diseases was of great value for further experimental studies.

#### Supplementary Note 6: Analysis of KDDANet resulting subnetworks mediating the associations between drugs and cancer using cancer-omics' datasets.

We tested whether KDDANet provides novel molecular insights on KDDAs mediating the associations between drugs and cancers using cancer-omics dataset. For this, we required that the type of cancer of KDDA was contained in TCGA, GSDC1000 or CCLE. KDDANet predicted 1847 novel unknown genes mediating KDDAs mediating the associations between drugs and cancer. We found that 149 of these genes were contained in COSMIC Cancer Gene Census with ~96-fold enrichment and adjusted *p*-value of 5.709e-10 (Hypergeometric test and Bonferroni correction). Using different types of TCGA datasets, including mRNA expression, Single Nucleotide Variant (SNV), Copy Number Variant (CNV) and DNA methylation, we checked whether these genes undergo oncogenic alterations in tumor samples. As expected, we found that these KDDA candidate genes have significantly higher SNV and CNV ratio in tumor samples than randomly selected genes (Supplementary Figure 4a). In consistent with this, the expression abundances and methylation levels of these candidates are significantly dysregulated in tumor samples than randomly selected genes, respectively (Supplementary Figure 4a). We further checked whether these genes undergo oncogenic alterations in cancer cell lines. In line with TCGA primary tumor data, we observed that they harbor more SNVs and CNVs in cancer cell lines than randomly selected genes (Supplementary Figure 4b).

We further asked whether the SNVs and CNVs of KDDA genes mediating the associations between drugs and cancers were correlated with the responses of cancer cell lines under anti-cancer drug treatment. For each gene of KDDA subnetwork mediating the association between drug and cancer, we evaluated the difference of drug’s half maximal inhibitory concentration (IC50) between cell lines harboring SNVs or CNVs on this gene (HSCs) and cell lines without SNVs and CNVs (WSCs) on this gene using Mann-Whitney U test. We required that the KDDA was contained in GSDC1000 or CCLE. We observed that the difference of IC50 values of anti-cancer drugs between HSCs and WSCs derived from KDDA genes mediating the associations between drugs and cancers was more significant than IC50 values of anti-cancer drugs between HSCs and WSCs derived from randomly selected genes (Supplementary Figure 4c). Such as,

KDDANet predicted, TP53, a well-known tumor suppression gene, mediating doxorubicin-bladder cancer association and topotecan-glioma association. The SNVs of bladder cancer cell line on TP53 was correlated with decreased IC50 values of doxorubicin (Supplementary Figure 4d). And the SNVs of glioma cancer cell line on TP53 was correlated with increased IC50 values of topotecan (Supplementary Figure 4d). A key lung cancer driver gene CCND1 was predicted mediating pyrimethamine-lung cancer association [41]. The CNVs of lung cancer cell lines on CCND1 was correlated with decreased IC50 values of pyrimethamine (Supplementary Figure 4d). CTNND2 encoded δ-Catenin which promoted tumorigenesis and metastasis of lung adenocarcinoma [42]. CTNND2 was inferred mediating paclitaxel and lung cancer association. The CNVs of lung cancer cell lines on CTNND2 was correlated with decreased IC50 values of paclitaxel (Supplementary Figure 4d). Based on these, we further tested whether the expression abundances and methylation levels of KDDA genes mediating the associations between drugs and cancerd were also associated with the responses of cancer cell lines under anti-cancer drug treatment. When compared with randomly selected genes, the expression abundances and methylation levels of KDDA genes mediating the associations between drugs and cancers were more obviously correlated with IC50 values of anti-cancer drugs in both GSDC1000 and CCLE cancer cell lines, respectively (Supplementary Figure 4e). For example, SUMO1 has been linked to tumorigenesis and invasion in gastric cancer [43]. KDDANet inferred SUMO1 mediating etoposide-gastric cancer association. The expression level of SUMO1 was negatively correlated to IC50 value of etoposide in gastric cancer cell lines (Supplementary Figure 4f). It has been reported that PML as potential treatment for pancreatic cancer [44]. KDDANet discovered PML mediating etoposide-pancreatic cancer association. The methylation level of PML gene was positively correlated to IC50 value of etoposide in pancreatic cancer cell line (Supplementary Figure 4f). E2 ubiquitin-conjugating enzymes have been reported involving in various tumor-promoting processes [45]. KDDANets predicted UBE2I mediating doxorubicin-gastric cancer. The methylation level of UBE2I gene was negatively correlated with IC50 value of doxorubicin (Supplementary Figure 4f). We have not found that these correlations are reported in the GSDC1000 and CCLE studies. Collectively, these outcomes indicated that KDDANet provided novel molecular insights into the molecular mechanism of anti-cancer drug response of cancer cell lines.

#### Supplementary Note 7: An online web server for KDDA decoding.

To help biomedical researchers to understand the pathogenesis of disease and promote drug development, we develop an online web server, http://www.kddanet.cn, for facilitating the researchers to explore the subnetwork of genes mediating KDDAs. Our website is divided into three parts: I) querying and browsing subnetwork of genes mediating individual KDDA (iKDDA); II) querying and browsing meta-subnetwork of shared genes mediating multiple KDDAs connecting multiple diseases to one drug (MDiODr) and III) querying and browsing meta-subnetwork of shared genes mediating multiple KDDAs connecting multiple drugs to one disease (MDrODi). For Part I, subnetworks of genes mediating 52878 KDDAs are provided for querying, visualizing, and downloading, respectively. For Part II, 12386 shared meta-subnetworks are contained in our web server. For Part III, 773 shared meta-subnetworks are contained in our web server. We design a bulk download interface and tutorials for facilitating user to explorer KDDANet resulting subnetworks. Overall, we provide a simply and easy-to-use web server for user to decode the hidden genes mediating KDDAs and thus help researchers to understand the pathogenesis of disease, accelerate drug repurposing and disease treatment. We next provided a more in-depth description with some figures generated from our web server and one example as step-by-step schematic illustration to explain the utility of KDDANet. For Part I, the iKDDA page aims to search gene subnetwork mediating the association between a given drug and a given disease. We provided an example of searching gene subnetwork mediating the association between heparin and obesity. For Part II, the MDiODr page aims to search the shared gene meta-subnetwork mediating multiple KDDAs between a given drug and a given disease type. We provided an example of searching gene subnetwork mediating multiple associations between bivalirudin and cancer. For Part III, the

MDiODr page aims to search the shared gene meta-subnetwork mediating multiple KDDAs between a given disease and all its related drugs. We provided an example of searching gene subnetwork mediating multiple associations between 13 drugs and GRACILE syndrome (Finnish lethal neonatal metabolic syndrome). Please see the detailed tutorials as follow and the Help section of our website for detailed description of the utility of an online web version of KDDANet tool.

#### Tutorials of KDDANet

### iKDDA page

This page aims to search gene subnetwork mediating the association between a given drug and a given disease.

##### iKDDA Search Box

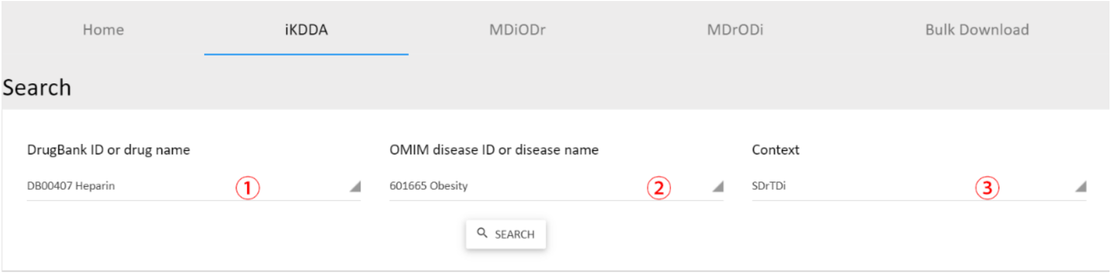


① Select drug from the drop-down list by DrugBank ID or drug name. A prompt will appear when you type characters.

② Select disease from the drop-down list by OMIM disease ID or disease name.

③ Select the context. SDrTDi: Network flow model constructed by using query drug as source node. SDiTDr: Network flow model constructed by using query disease as source node.

### iKDDA Search Result

#### iKDDA Subnetwork

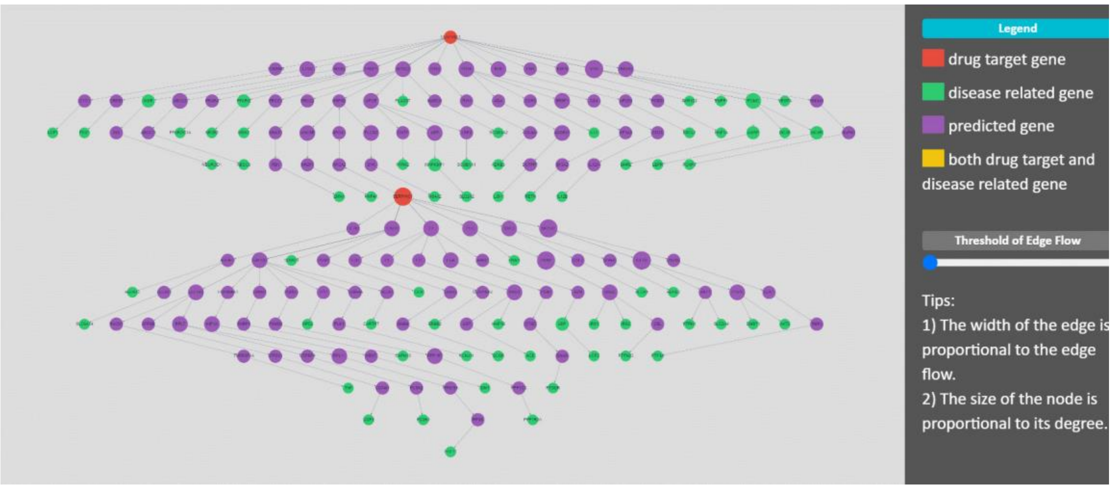


Tips:

1. Move the threshold of Threshold of Edge Flow slider in the right panel, it will dynamically display the updated gene subnetwork by selecting the edge with flow amount larger than the threshold. Bar left: minimal edge flow. Bar right: maximum edge flow.

2. Box selection via modifier key (shift, command, control, alt)

3. Mouse wheel to zoom

4. Grab and drag background to pan

5. Click a gene node to see the NCBI gene information

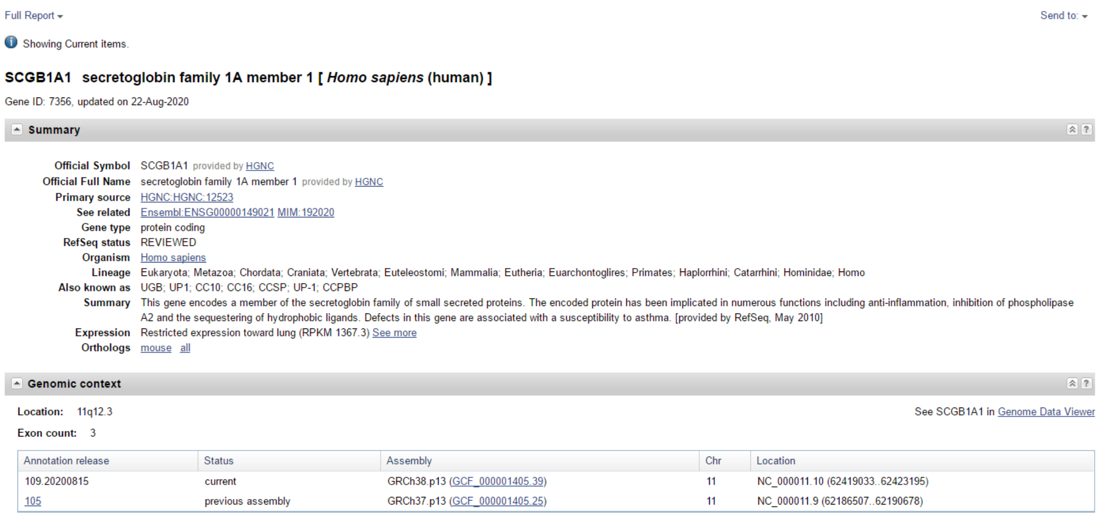

Full Report ▾ | Send to: ▾

Showing Current items.

**SCGB1A1 secretoglobin family 1A member 1 [ *Homo sapiens* (human) ]**

Gene ID: 7356, updated on 22-Aug-2020

**Summary**

| | |
|---|---|
| **Official Symbol** | SCGB1A1 provided by HGNC |
| **Official Full Name** | secretoglobin family 1A member 1 provided by HGNC |
| **Primary source** | HGNC:HGNC:12523 |
| **See related** | Ensembl:ENSG00000149021 MIM:192020 |
| **Gene type** | protein coding |
| **RefSeq status** | REVIEWED |
| **Organism** | Homo sapiens |
| **Lineage** | Eukaryota; Metazoa; Chordata; Craniata; Vertebrata; Euteleostomi; Mammalia; Eutheria; Euarchontoglires; Primates; Haplorrhini; Catarrhini; Hominidae; Homo |
| **Also known as** | UGB; UP1; CC10; CC16; CCSP; UP-1; CCPBP |
| **Summary** | This gene encodes a member of the secretoglobin family of small secreted proteins. The encoded protein has been implicated in numerous functions including anti-inflammation, inhibition of phospholipase A2 and the sequestering of hydrophobic ligands. Defects in this gene are associated with a susceptibility to asthma. [provided by RefSeq, May 2010] |
| **Expression** | Restricted expression toward lung (RPKM 1367.3) See more |
| **Orthologs** | mouse all |

**Genomic context**

**Location:** 11q12.3 | See SCGB1A1 in Genome Data Viewer

**Exon count:** 3

| Annotation release | Status | Assembly | Chr | Location |
|---|---|---|---|---|
| 109.20200815 | current | GRCh38.p13 (GCF_000001405.39) | 11 | NC_000011.10 (62419033..62423195) |
| 105 | previous assembly | GRCh37.p13 (GCF_000001405.25) | 11 | NC_000011.9 (62186507..62190678) |

### Subnetwork Download

DOWNLOAD

DB00407_601665_download.txt

#### File Content

| | | | | |
|---|---|---|---|---|
| 3339 | HSPG2 | 344 | APOC2 | 0.00746269 |
| 3339 | HSPG2 | 335 | APOA1 | 0.0223881 |
| 6514 | SLC2A2 | 9479 | MAPK8IP1 | 0.0149254 |
| 2206 | MS4A2 | 6850 | SYK | 0.0149254 |
| 3596 | IL13 | 1232 | CCR3 | 0.0149254 |
| 5506 | PPP1R3A | 5501 | PPP1CC | 0.0149254 |
| 10554 | NA | 1967 | EIF2B1 | 0.0298507 |
| 3827 | KNG1 | 2160 | F11 | 0.0298507 |
| 7356 | SCGB1A1 | 4036 | LRP2 | 0.0149254 |
| 56729 | RETN | 8175 | SF3A2 | 0.0149254 |

Column 1: Gene 1 EntrezID

Column 2: Gene 1 Name

Column 3: Gene 2 EntrezID

Column 4: Gene 2 Name

Column 5: Edge Flow

### MDiODr page

This page aims to search the shared gene meta-subnetwork mediating multiple KDDAs between a given drug and a given disease type.

#### MDiODr Search Box

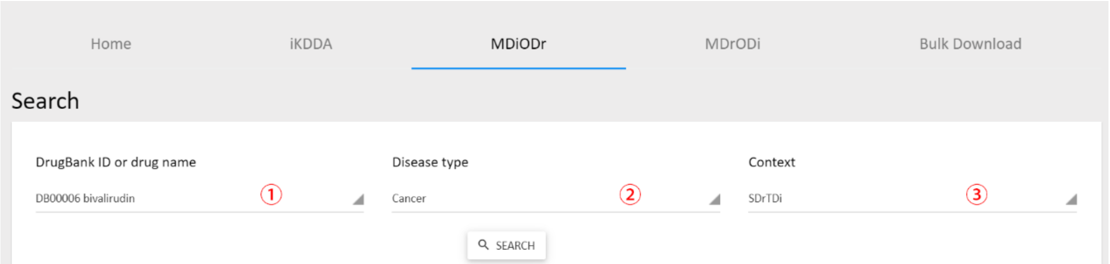


① Select drug from the drop-down list by DrugBank ID or drug name. A prompt will appear when you type characters.

② Select disease type from the drop-down.

③ Select the Context. SDrTDi: Network flow model constructed by using query drug as source node. SDiTDr: Network flow model constructed by using query disease as source node.

#### MDiODr Search Result

#### Related Diseases

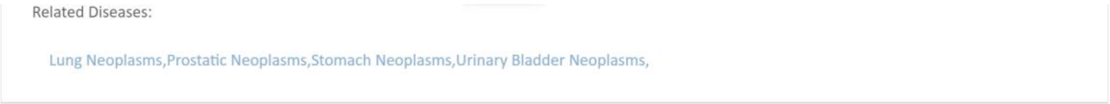


Click a disease to see the OMIM disease information

# 211980

LUNG CANCER

Other entities represented in this entry:

ALVEOLAR CELL CARCINOMA, INCLUDED
ADENOCARCINOMA OF LUNG, INCLUDED
NONSMALL CELL LUNG CANCER, INCLUDED
LUNG CANCER, PROTECTION AGAINST, INCLUDED

**Phenotype-Gene Relationships**

| Location | Phenotype | Phenotype MIM number | Inheritance | Phenotype mapping key | Gene/Locus | Gene/Locus MIM number |
|---|---|---|---|---|---|---|
| 1q24.3 | {Lung cancer, susceptibility to} | 211980 | AD, SMu | 3 | FASLG | 134638 |
| 2q33.1 | {Lung cancer, protection against} | 211980 | AD, SMu | 3 | CASP8 | 601763 |

#### MDiODr Meta-Subnetwork

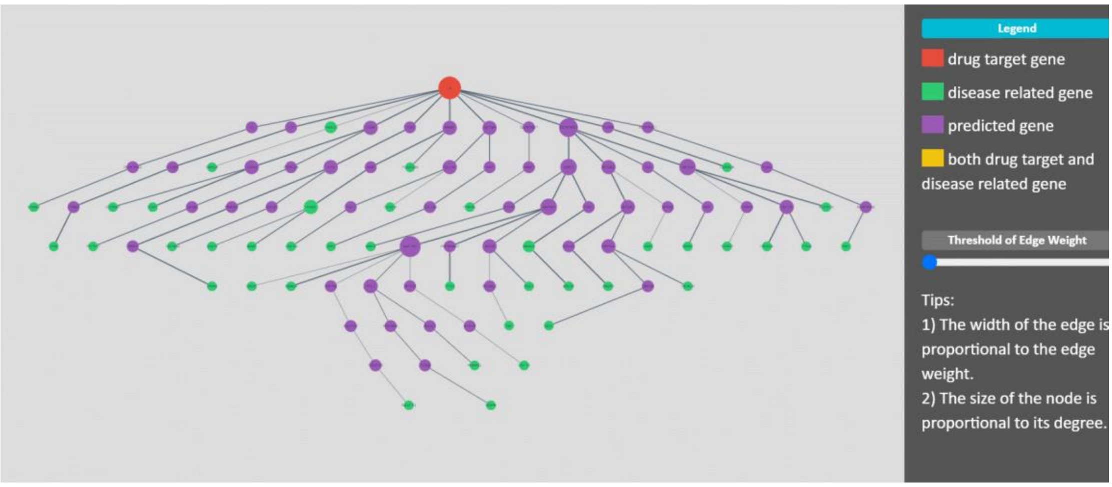


Tips:

1. Move the threshold of Threshold of Edge Weight slider in the right panel, it will dynamically display the updated gene subnetwork by selecting the edge with weight larger than the threshold. Bar left: minimal edge weight. Bar right: maximum edge weight.

2. Box selection via modifier key (shift, command, control, alt)

3. Mouse wheel to zoom

4. Grab and drag background to pan

5. Click a gene node to see the NCBI gene information

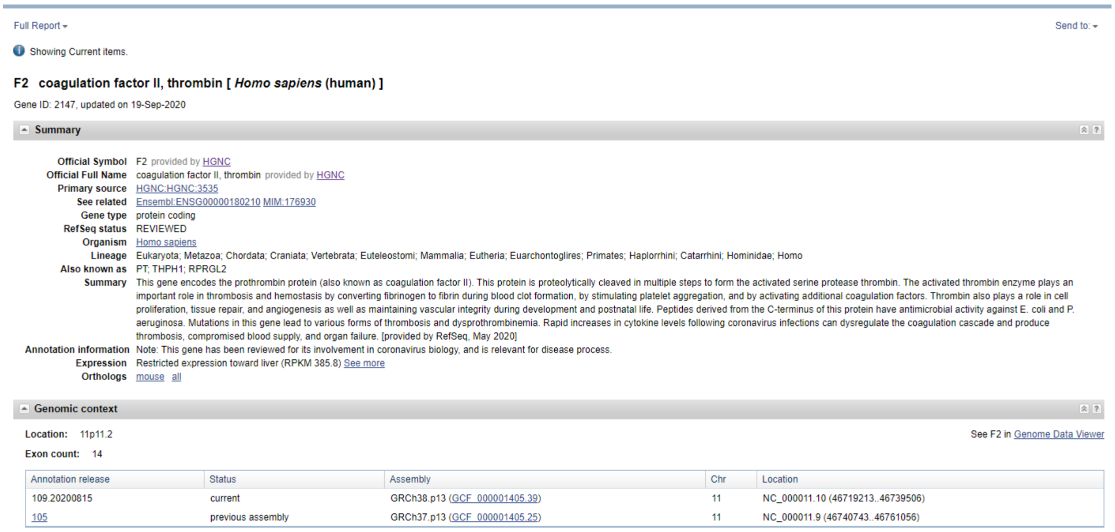


### Download

DOWNLOAD

DB00006_Cancer_download.txt

#### File Content

| | | | | |
|---|---|---|---|---|
| 6041 | RNASEL | 6059 | ABCE1 | 0.5 |
| 6129 | RPL7 | 1915 | EEF1A1 | 0.5 |
| 60 | ACTB | 2597 | GAPDH | 0.75 |
| 335 | APOA1 | 336 | APOA2 | 1 |
| 5781 | PTPN11 | 2885 | GRB2 | 0.75 |
| 1621 | DBH | 338 | APOB | 0.25 |
| 5111 | PCNA | 121504 | HIST4H4 | 0.5 |
| 2885 | GRB2 | 7916 | PRRC2A | 0.75 |

Column 1: Gene 1 EntrezID

Column 2: Gene 1 name

Column 3: Gene 2 EntrezID

Column 4: Gene 2 Name

Column 5: Edge Weight

### MDrODi page

This page aims to search the shared gene meta-subnetwork mediating multiple KDDAs between a given disease and all its related drugs.

#### MDrODi Search Box

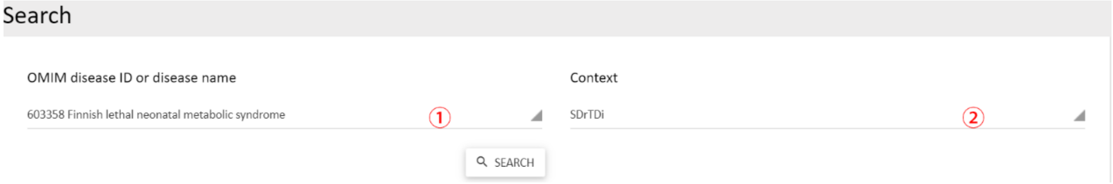


① Select disease from the drop-down list by OMIM disease ID or disease name. A prompt will appear when you type characters.

② Select the Context. SDrTDi: Network flow model constructed by using query drug as source node. SDiTDr: Network flow model constructed by using query disease as source node.

#### MDrODi Search Result

##### Related Drugs

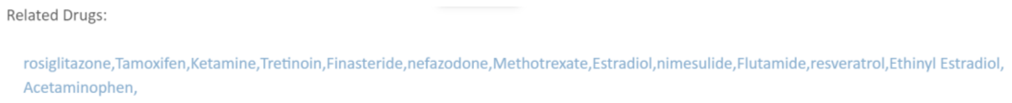


Click a drug to see the DrugBank drug information

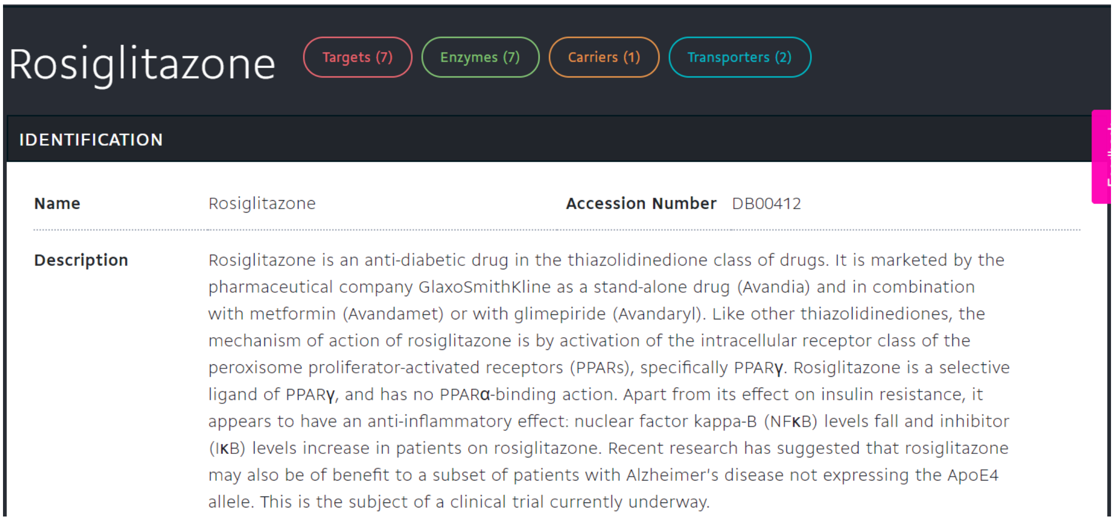


### MDrODi Meta-Subnetwork

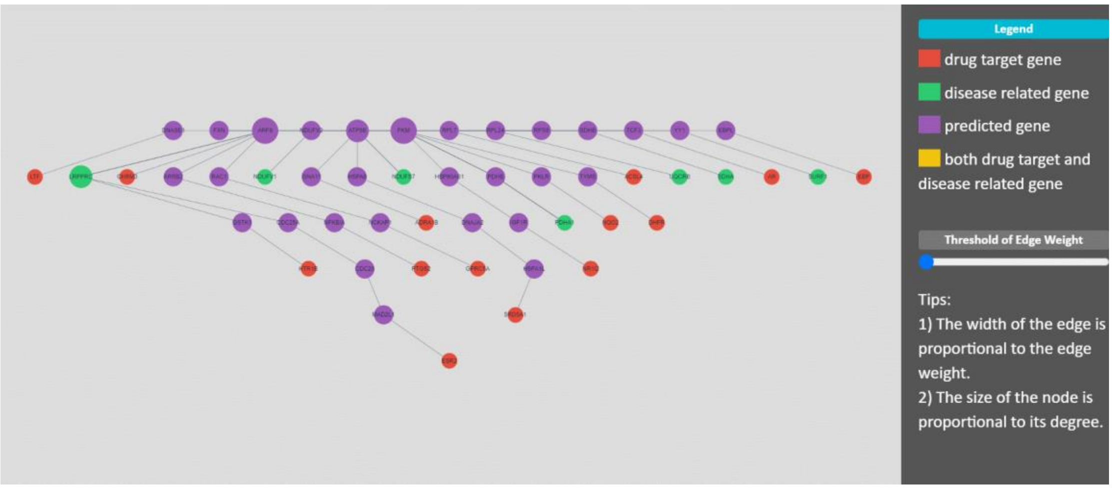


Tips:

1. Move the threshold of Threshold of Edge Weight slider in the right panel, it will dynamically display the updated gene subnetwork by selecting the edge with weight larger than the threshold. Bar left: minimal edge weight. Bar right: maximum edge weight.

2. Box selection via modifier key (shift, command, control, alt)

3. Mouse wheel to zoom

4. Grab and drag background to pan

5. Click a gene node to see the NCBI gene information

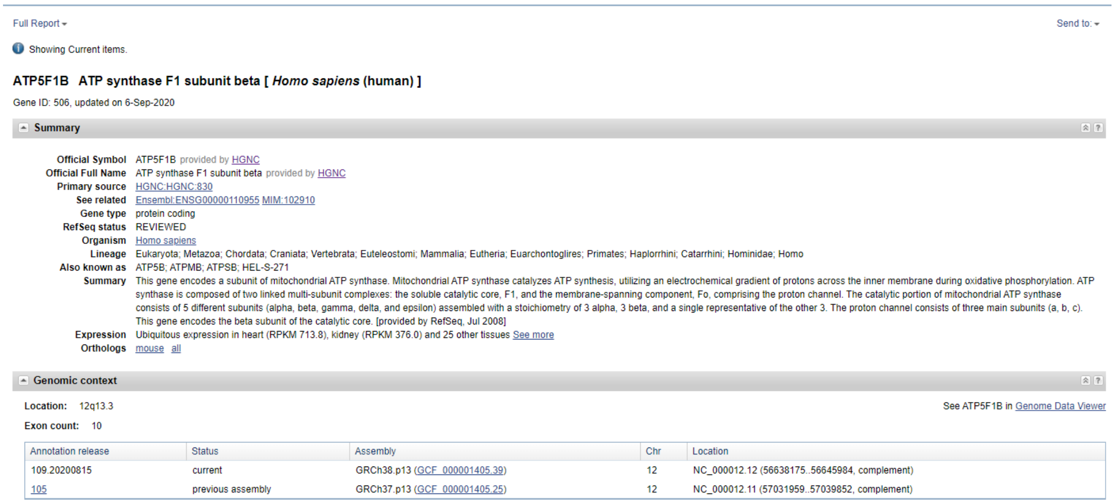

Full Report ▾ Send to: ▾

Showing Current items.

**ATP5F1B ATP synthase F1 subunit beta [ *Homo sapiens* (human) ]**

Gene ID: 506, updated on 6-Sep-2020

**Summary**

| | |
|---|---|
| **Official Symbol** | ATP5F1B provided by HGNC |
| **Official Full Name** | ATP synthase F1 subunit beta provided by HGNC |
| **Primary source** | HGNC:HGNC:830 |
| **See related** | Ensembl:ENSG00000110955 MIM:102910 |
| **Gene type** | protein coding |
| **RefSeq status** | REVIEWED |
| **Organism** | Homo sapiens |
| **Lineage** | Eukaryota; Metazoa; Chordata; Craniata; Vertebrata; Euteleostomi; Mammalia; Eutheria; Euarchontoglires; Primates; Haplorrhini; Catarrhini; Hominidae; Homo |
| **Also known as** | ATP5B; ATPMB; ATPSB; HEL-S-271 |
| **Summary** | This gene encodes a subunit of mitochondrial ATP synthase. Mitochondrial ATP synthase catalyzes ATP synthesis, utilizing an electrochemical gradient of protons across the inner membrane during oxidative phosphorylation. ATP synthase is composed of two linked multi-subunit complexes: the soluble catalytic core, F1, and the membrane-spanning component, Fo, comprising the proton channel. The catalytic portion of mitochondrial ATP synthase consists of 5 different subunits (alpha, beta, gamma, delta, and epsilon) assembled with a stoichiometry of 3 alpha, 3 beta, and a single representative of the other 3. The proton channel consists of three main subunits (a, b, c). This gene encodes the beta subunit of the catalytic core. [provided by RefSeq, Jul 2008] |
| **Expression** | Ubiquitous expression in heart (RPKM 713.8), kidney (RPKM 376.0) and 25 other tissues See more |
| **Orthologs** | mouse all |

**Genomic context**

**Location:** 12q13.3 See ATP5F1B in Genome Data Viewer

**Exon count:** 10

| Annotation release | Status | Assembly | Chr | Location |
|---|---|---|---|---|
| 109.20200815 | current | GRCh38.p13 (GCF_000001405.39) | 12 | NC_000012.12 (56638175..56645984, complement) |
| 105 | previous assembly | GRCh37.p13 (GCF_000001405.25) | 12 | NC_000012.11 (57031959..57039852, complement) |

### Download

#### File Content

| | | | | |
|---|---|---|---|---|
| 374291 | NDUFS7 | 506 | ATP5B | 0.153846153846154 |
| 2767 | GNA11 | 147 | ADRA1B | 0.0769230769230769 |
| 3305 | HSPA1L | 6715 | SRD5A1 | 0.0769230769230769 |
| 382 | ARF6 | 409 | ARRB2 | 0.0769230769230769 |
| 6152 | RPL24 | 6202 | RPS8 | 0.0769230769230769 |
| 5315 | PKM | 3326 | HSP90AB1 | 0.0769230769230769 |
| 5162 | PDHB | 5315 | PKM | 0.230769230769231 |
| 7528 | NA | 1773 | DNASE1 | 0.0769230769230769 |

Column 1: Gene 1 EntrezID

Column 2: Gene 1 name

Column 3: Gene 2 EntrezID

Column 4: Gene 2 Name

Column 5: Edge Weight

### Bulk Download page

This page aims to download all KDDA subnetworks for downstream analysis.

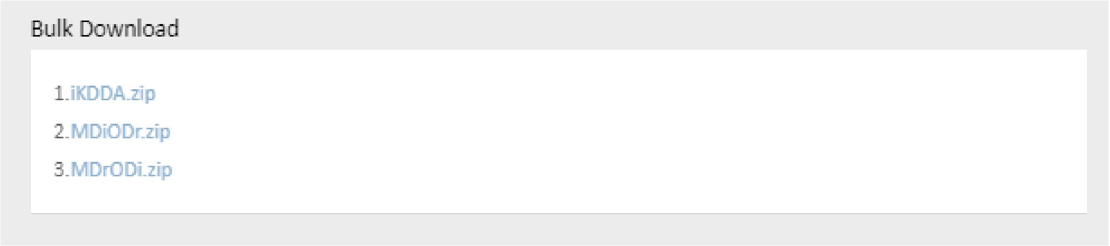
